# In vitro models to mimic tumor endothelial cell-mediated immune cell reprogramming in lung adenocarcinoma

**DOI:** 10.1101/2024.10.13.618092

**Authors:** Morgane Krejbich, Judith Fresquet, Emilie Navarro, Virginie Forest, Marine Cotinat, Valentin Isen, Hortense Perdrieau, Camille Chatelain, Vincent Dochez, David Roulois, Nicolas Boisgerault, Richard Redon, Isabelle Corre, Christophe Blanquart, Lucas Treps

## Abstract

Tumor endothelial (TECs) cells play a critical role in regulating immune responses within the tumor microenvironment (TME). However, the mechanisms by which TECs modulate immune cell population remain unclear, particularly in non-small cell lung cancer (NSCLC). Here, we investigated how NSCLC cells tweak normal endothelial cells (NECs) into TECs and the subsequent effects on immune regulation. NECs were cocultured with various NSCLC cell lines, using 2D and 3D coculture models to evaluate TEC-mediated effects on immune cells. We show that direct coculture led to significant transcriptomic, proteomic and kinomic alterations in TECs, especially in pro-inflammatory pathways. We identified a downregulation of the co-stimulatory molecule OX40L in TECs compared to NECs, suggesting impaired T-cell proliferation support. While TECs showed a limited effect on CD8+ T-cell activation, TECs supported CD4 T-cells polarization into Treg and Th22 subsets. Moreover, TECs also promoted M2-like macrophages polarization, thereby potentially contributing to the TME immunosuppression. State-of-the-art single-cell RNA sequencing of 3D multicellular tumor spheroids (MCTS) revealed distinct TEC subpopulations, including an inflammatory subset with UPR signature. The latter was absent in 2D-cultured NECs but present in freshly isolated and 2D-cultured TECs from NSCLC patients. Importantly, we also identified within MCTS a perivascular M2-like macrophage subset, predicted to interact with TECs with MIF and Midkine signaling. In conclusion, TECs in NSCLC tumors play a pivotal role in remodeling the TME immune landscape by promoting immune suppression. This study highlights the complex immunoregulatory functions of TECs within our different *in vitro* models that mimic aspects of the TME. Our data may provide new insights into potential therapeutic strategies targeting TECs or regulatory signaling to improve the efficacy of immunotherapy in NSCLC.

## Introduction

Responsible for an estimated 1.8 million deaths per year, lung cancer is one of the leading causes of cancer incidence and mortality worldwide, with non-small-cell lung cancers (NSCLC) accounting for 80-85% of cases ^1^. Over the past decade, a better understanding of the biology of lung cancer, particularly the composition and activity of the tumor microenvironment (TME), has led to the development of new therapies that have improved survival ^2^. As such, immunotherapy in the form of immune checkpoint inhibitors have revolutionized patient outcome compared to chemotherapy alone ^3–6^.

These immunotherapies are based on the reversal of the anergy of T lymphocytes that is induced in the TME by tumor cells themselves, heterogeneous immune cells including tumor-associated macrophages (TAM), regulatory T cells (Treg) and Th2/22 cells. However, tumor progression and response to treatment may also be affected by structural cells, including cancer-associated fibroblasts (CAFs), endothelial cells (ECs), and other factors such as extracellular matrix. There is considerable evidence that tumor endothelial cells (TECs) have unique phenotypic and functional traits in comparison to normal endothelial cells (NECs) in terms of metabolism, genetics and transcriptomic profile ^7^. Moreover, a recent set of evidence indicates that TECs display immunoregulatory features that could be therapeutically relevant for cancer patients and their response to immunotherapies ^8–13^. Current assumptions are that some TECs subsets, in the form of tumor-associated high endothelial venules (HEVs) are gatekeeper for TME infiltrating immune cells and essential for successful antitumor immunity ^14^, whereas other TECs act as semi-professional antigen presenting cells (APC) to activate or inhibit effector cells ^15^ and model tertiary lymphoid structures in the tumor ^16^. The difficulty in studying TECs holds in the fact that they are highly heterogeneous and plastic by nature and the lack of relevant models to study their immunoregulatory characteristics. In this study we established various models to unravel how tumor cells could remodel NECs into TECs, and how their remodeling could impact the immune system. Various approaches including single-cell RNA sequencing (scRNA-seq) and functional assays are employed to reach this aim.

## Materials & methods

### HUVEC isolation

Human umbilical vein endothelial cells (HUVECs) were freshly isolated from umbilical cords obtained from multiple donors from the CHU Maternity, Nantes. Informed consent was obtained from all subjects. The blood from umbilical cord is rinsed with PBS and sterile gauze and 10 ml of the 0,2% collagenase type I in 0,9% NaCl + 2 mM CaCl2 + 2X P/S, PBS and M199 solution is added in the vein and the up side clamped with a sterile clip. The umbilical cord is incubated 13 min at 37 degrees. After incubation, the umbilical cords are unclipped and the collagenase collected. The vein is rinsed with M199 medium and the cells are centrifuged at 1200 rpm for 5 minutes. The cells are resuspended into a M199 medium with 2X P/S and grown on a gelatin coated T75 flask in a 37 degrees incubator with 5% of CO2.

### Cell culture, Coculture and CD31 sorting

HUVECs were cultured in Endothelial Cell Growth Medium 2 (EGM2) media (22011B, Promocell) supplemented with 2% fetal calf serum (FCS), 5 ng/ml epidermal growth factor (EGF), 10 ng/ml basic fibroblast growth factor, 20 ng/ml Insulin-like Growth Factor (Long R3 IGF), 0.5 ng/ml VEGF 165, 1µg/ml ascorbic acid, 22.3 µg/ml heparin, and 0.2 µg/ml hydrocortisone from the Supplement Mix (C-39216) in flasks coated with 0.1% gelatin

(Sigma; Lot#SLCF9893). Cells were used between passages 2-7. NSCLC cell lines, A549, H1755 and H1975 were cultured in RPMI 1640 medium (Gibco (Invitrogen)) supplemented with 100 U/ml penicillin, 100 mg/ml streptomycin, 2 mM L-glutamine (Gibco (Invitrogen)), 10% FBS (Corning), and cultured at 37°C in a 5% CO2 atmosphere. All cells were tested each week to prevent mycoplasma contaminations using PlasmoTest™ (Invivogen).

We conducted our coculture according to the protocol suggested by Njock and his team ^17^. The day before coculture HUVECs and NSCLC cell lines were rinsed with D-PBS and placed in EGM2 without growth factor (EGM2 SF) but completed with 0.5% FBS exofree (extracellular vesicles depleted) and 5 ng/ml human Fibroblast Growth Factor (bFGF) (HY- P7004, Med Chem Express) overnight. The next day, HUVECs were seeded in EGM2 for 6 hours, then tumor cells were added to plates with adherent HUVECs in a 1:1 ratio in EGM2 SF, 0.5% FBS exofree and 5 ng/ml human bFGF. After 48 hours, monoculture (HUVEC alone) or coculture (HUVEC+A549/ HUVEC+H1755/ HUVEC+H1975) are detached with TrypLE Select (1X) (Gibco^TM^,12563029) and HUVEC were positively selected for CD31 by magnetic cell sorting (MACS) using LS separation columns (Miltenyi Biotech, 130-042-401). Following the manufacturer’s instructions, a positive staining of HUVEC using the CD31 MicroBead Kit (Miltenyi Biotech, 130-091-935) was performed. Cells were passed sequentially over two separation columns to reach higher purity. EGM2 was used for the final elution step from the column.

### Monocyte and Lymphocyte T-Cell Purification

Peripheral Blood Mononuclear cells (PBMCs) were collected from healthy donors (Etablissement Français du Sang, ethics agreement CPDL-PLER-20** **), isolated using Ficoll gradient (Eurobio, Cat#CMSMSL01-01). Polyclonal CD8+ T cells were selected using EasySep Human CD8+ T Cell Isolation Kit (STEMCELL Technologies; 17953). Blood monocytes were isolated through negative magnetic sorting (EasySep Human Monocyte Enrichment Kit, StemCell)

### Myeloid cell differentiation

Macrophages were differentiated from monocytes over 4C:days, seeded in a 12-well plate at 1.10^6^ cells in 1C:mL of RPMI supplemented as described. At the seeding day and the day after, 200C:U/mL of GM-CSF (Miltenyi) was added to differentiate M1-like macrophages or 5000C:U/mL of M-CSF (Miltenyi) for M2-like macrophages. After 4C:days, macrophages were harvested for experiments.

### RNA isolation

After CD31 sorting, total RNA from HUVECs was extracted using the miRNeasy Micro Kit (Qiagen; 217084) according to the manufacturer’s instruction, and RNA integrity and profiles were measured before sequencing using the Agilent 2100 Bioanalyzer (Agilent) for total RNA (RNA nanochips).

### Reverse-transcriptase quantitative polymerase chain reaction (RT-qPCR)

One microgram of RNA was reverse transcribed using Revert Aid H Minus Reverse Transcriptase (Thermo Scientific), and the RT product was used for expression analysis using Maxima SYBR Green/ROX qPCR Master Mix (Thermo Scientific) on a QuantStudio3 system (Applied). Targets of interest were amplified using premade primer: PECAM (QT00081172), ICAM (QT00074900), VCAM (QT00018347), PD-L1 (QT00082775), TNFSF4 (QT00028658). Gene expression was normalized based on the expression of the housekeeping gene encoding the Ribosomal Protein Lateral Stalk Subunit P0 (RPLP0).

### 3’RNA-seq digital gene expression and analysis

Gene analysis was performed thanks to the GenoBird platform by 3’RNA-sequencing profiling using a NovaSeq 6000 (Illumina). The quality of raw sequence reads was assessed by FastQC. Adapter sequences were trimmed off the raw sequence reads using Cutadapt. Reads were aligned to the human (hg38) genome using BWA. All obtained data have been uploaded on GEO Omnibus site (GSE247526). Bioinformatic analysis to compare gene expression signature and gene set enrichment analysis was carried out using the DESeq2 and pathfindR packages. An adjusted p-value <0.05 was considered statistically significant.

### MCTS formation and scRNA-seq

MCTS were formed by mixing A549, normal primary HUVECs, normal primary human lung fibroblast, and PBMC-derived monocytes at a defined ratio of 2:1:1:1, with a total of 20.000 cells per MCTS. Cells were mixed and loaded in a low adherence plate, and cells pelleted by centrifugation. The EGM2 medium was used to culture the MCTS.

After 5 days of culture, MCTS were collected, rinsed with PBS and dissociated with TrypLE. The single cell suspension was then resuspended in a 0.04% BSA and viability assessed.

33.000 cells were used to generate single cell droplet librairies with Chromium NEXT GEM Single Cell 3’ Reagent kit v3.1, 10X Chromium Single Cell Controller (10X Genomics, Pleasanton, CA, USA). Library size was determined by Agilent TapeStation assays and the concentration by Qubit Flex (Invitrogen). Libraries were pooled and sequenced on NovaSeq 6000 Sequencing System (Illumina), providing a read depth of >20,000 read pairs per cell according to manufacturer’s instructions. A number of 4 samples was sequenced, representing different donors of fibroblasts, HUVECs and PBMC-derived monocytes. The A549 was the same across all 4 samples and was used with no more than 2 passages in between the 2 batches.

### scRNA-seq bioinformatic analysis

Gene expression matrices were generated using Cell Ranger (10X Genomics, version 2.1.1) using the GRCh38 build of the human reference genome, and further processed using RStudio (version 2023.12.1+402). We used the following quality control steps: genes expressed by <3 cells were not considered, cells expressing <200 genes (low quality) or >10.000 genes, <200 genes or >10.000 unique molecular identifiers (UMIs), or >10% of UMIs derived from the mitochondrial genome were removed. 279 doublets were detected and removed using the DoubletFinder package. Doublets represent ∼0.5% of filtered cells and did not form any specific cluster nor were overrepresented in a particular sample. Visualization, clustering, marker gene identification and gene set enrichment analysis of the remaining cell data was performed using the SeuratV5 package. Prior to <u>SCENIC analysis</u>, gene expression matrix was exported to python compatible file using ScopeLoomR. Then, after 10 iterations of pySCENIC pipeline, results were aggregated in custom python pipeline. Using only matrices resulting from the aggregation, the integration of regulons data was analyzed in Seurat and R. <u>Copy number variations</u> (CNV) were estimated using the *inferCNV* R package (version 1.5.0). For this analysis, we first performed the analysis without reference populations to visualize chromosomal alterations across all cell types. After identifying cluster with similar CNV, and matching these results with cluster gene signatures, we rerun inferCNV with the immune cell, ECs and fibroblasts as reference cells. Analysis was performed using the following parameters: cut-off 0.1, denoise = T, min_cells_per_gene = 10, HMM = F, cluster_by_groups = F. <u>CellChat</u> was performed using default parameters ^18^ on the full CellChat database by predicting interactions between the endothelial and myeloid compartments. A <u>meta-analysis</u> for commonly upregulated gene sets between HUVEC monocultures and NSCLC or MDA-MB-231 cocultures ^17^ was performed using the BIOMEX software ^19^.

### Protein Tyrosine Kinase (PTK) and Serine Threonine Kinase (STK) kinome assay

After NSCLC-HUVECs isolation, cell pellets are collected and lysed with a M-PER TM reagent (Thermo Scientific, 78501) containing protease and phosphatase inhibitors (Thermo Scientific, 87785). Lysates were dosed using BCA to make sure that we had enough protein quantity for all our experimental conditions and 3.35µg and 1µg of protein samples were loaded on the PTK and STK PamChip arrays, respectively (PamGene). Everything was processed using the manufacturer’s recommendations and using the PamStation 12. The array allows the tracking of the phosphorylation of 196 peptides for the PTK and of 144 peptides for the STK, where the kinase activities are predicted, and further checked for quality filtering. The data were analyzed by PamGene and the pathways identified with the EnRichr website.

### Secretome proteomic analysis

Digestions and LC-MS/MS analyses were performed at the Prot’ICO proteomics facility. NSCLC cell lines were incubated 72 hours in RPMI with 10%SVFd and 1% PS. Supernatants were washed away with 1X PBS and replaced by RPMI without SVF and allowed to condition this new medium for 24 hours. Cells were counted for subsequent normalization, and media (∼6 mL) were collected and centrifuged 5 min at 300*g* and concentrated on an amicon spin reverse 5 kDa/2mL membrane filtration device. Proteins were denatured in 0.1% Rapigest SF™ acid-labile detergent (Waters^®^), 5 mM DTT and 50 mM ammonium bicarbonate, at 95°C for 30 min (200 µL final). Thiol residues were thus chemically reduced and subsequently cooled down to RT then protected by alkylation in 20 mM MMTS (Sigma Aldrich^®^) for 10 min at 37°C. 5µg of trypsin (ABSciex) per sample were added and incubated at 37°C overnight. Peptides were then cleared by centrifugation, desalted on C_18_ sep-pack reverse phase microcolumns as described in the “Stage-tips” procedure and peptides were eluted in Acetonitrile (ACN)^23^. Eluates were dried in a vacuum centrifuge concentrator (Thermo), resuspended in 25 µL of 10% ACN and 0.1% Formic Acid (FA). Eluate flow was electrosprayed into a timsTOF Pro 2 mass spectrometer (also from Bruker®) for the 60 min duration of the hydrophobicity gradient ranging from 99% of solvent A (0.1% FA in milliQ-grade H_2_O) to 40% of solvent B (80% ACN and 0.1% FA in mQ-H_2_O). The mass spectrometer acquired data throughout the elution process and operated in data-independent analysis mode (DIA) with PASEF-enabled method using the TIMS-Control v.3.1.4 software (Bruker^®^). Samples were injected in batch replicate order to circumvent possible technical biases.

LC-MS/MS data analysis: The raw data were extracted, normalized and analyzed using Spectronaut software v. 18.6.231227.55695 (Biognosys®) in Direct-DIA mode, which modelized elution behavior, mobility and MS/MS events based on the Uniprot/Swissprot sequence 2022 database of human proteins. Protein identification false discovery rate (FDR) was restricted to 1% maximum, with match between runs (MBR) option enabled and inter-injection data normalization.

### Flow cytometry

To characterize Endothelial cell after sorting:

Cells were stained in suspension for 30 min at 4°C with anti-CD31-AlexaFluor 488 (Biolegend, 303110), anti-CD274-PE (BD, 557924), anti-CD252-PE (Biolegend, 326308), anti-CD54-APC-Fire 750 (Biolegend, 353122), anti-CD106-APC (Biolegend, 305810). Cells were washed twice with PBS, processed by flow cytometry on a 5 laser, 32 detector BD FACSymphony™ A5 Cell Analyzer (BD Bioscience) (Acquisition software: Diva 8) and analyzed with the FlowJo VX software. Median fluorescence intensities measured in flow cytometry experiments are normalized on the monoculture conditions and are displayed as ratios of median fluorescence intensities (RMFI) ± standard error of the mean (SEM).

To characterize macrophage populations after 5 days cocultured with sorted HUVECs:

After washing in Phosphate buffered saline (PBS), the cells were incubated 20 min with 1:500 Zombie UV™ Fixable Viability dye (Biolegend) in PBS. They were washed with PBS- 0.1% Bovine Serum Albumine (BSA, Sigma-Aldrich) and fixed with PBS-4% paraformaldehyde (PFA, Electron Microscopy Sciences) for 10 min, washed and stored at 4°C.

Cells were incubated in 50C:µL of Brilliant Stain Buffer (BD Biosciences) supplemented with 1.25C:µg of human BD Fc Block (BD Biosciences). After 10 min, appropriate concentrations of specific antibodies were added for 30C:min at 4°C with 1µg/ml anti-CD31-AlexaFluor 488 (Biolegend, 303110), 1:800 anti-CD45-BUV805 (BD Biosciences), 1:200 anti-CD163-BV711 (BD Biosciences), 1:400 anti-HLA-DR-PE Vio770 (Miltenyi), 1µg/ml anti-CD16-BUV737 (BD Biosciences) and 1:100 anti-CD14-APC Vio770 (Miltenyi). After three washes in PBS- 0.1%BSA, stained cells were acquired on a 5 laser, 32 detector BD FACSymphony™ A5 Cell Analyzer (BD Bioscience). The BD FACSDiva 8.0 software (BD Biosciences) was used for data acquisition and analyzed with the FlowJo VX software.

### Confocal Microscopy

Cocultures were seeded into 24-well plates on coverslips for 48 hours and then fixed in PBS- 4% PFA during 20 minutes at room temperature (RT). PBS-2%Triton-1%BSA was used to permeabilize the cells for 30min at RT. Primary antibodies anti-CD31-488 (Biolegend, 303110) anti-vWF (Abcam, ab6994), CD163-594 (Biolegend, 364304), CD45-647 (Biolegend, 304018) were incubated with the cells overnight at 4°C, and secondary antibodies AlexaFluor 594 (Cell Signalling Technology, 8889S) were incubated for 1 hour at RT with Hoechst 33342 (Sigma, 14533) and anti-Phalloidin-647 (Invitrogen, A30107). Pictures were taken with a Nikon A1rHD LFOV confocal microscope, with a 60x oil immersion objective (Nikon Instruments). The images were processed and analyzed using the ImageJ software.

After formation, MCTS spheroids were collected in PBS and permeabilized in PBS Triton 2%. After primary and secondary antibody incubations, MCTS were rinsed and immobilized on a IBIDI 8p with Cell-Tak (Corning). Finally, MCTS are transparized using RapiClear (SunjinLab, Nikon) 1.47 before image acquisition.

### ELISA assay

Cytokine production was quantified from cocultured supernatants 24 hours after exposing LTs to HUVECs. TNF-α secretion was measured according to the manufacturer’s instructions using uncoated TNF-α ELISA kits (Invitrogen, 88-7346-88) and absorbance values were read at 450/570nm using Multiskan FC microplate reader (Thermo Scientific) and concentrations were calculated according to the manufacturer’s instructions. IL-10 secretion was measured according to the manufacturer’s instructions using uncoated IL-10 ELISA kits (Biolegend, 430604) and absorbance values were read at 450/570nm using Multiskan FC microplate reader (Thermo Scientific). IL-10 concentrations were calculated according to the manufacturer’s instructions.

### Chemokine detection by multiplex assay

The supernatants of cocultures were collected after 5 days of coculture between sorted HUVEC and naïve CD4+ T Lymphocytes. Cytokines such as IL-5, IL-13, IL-2, IL-6, IL-9, IL- 10, IFN-γ, TNF-α (TNFSF2), IL-17A, IL-17F, IL-4, IL-22 were quantified using the LEGENDplex™ Human T Helper Cytokine Panels Version 2 (BioLegend) according to the manufacturer’s recommendations.

### Leukocytes Adhesion assay

After CD31 sorting, HUVEC were grown in 24-well plates coated with gelatin to obtain a confluent cell monolayer during 6 or 24 hours. Over this period, the activated monoculture control is exposed to cytokine cocktail (6nM TNF-α and IL1-β). PBMCs were obtained as described above and labelled with calcein AM (Life, C3100MP). Medium was removed and HUVECs were washed with PBS before the addition of 2.0 x 10^6^ and incubation for 1 hour at 37°C. Non-adherent cells were removed by delicate washing 7 times with PBS. Cellular adhesion rates were determined by live cell imaging using the IncuCyte (Sartorius). Around ten fields-of-view per well are randomly chosen and analyzed with Fiji ^20^ to determine the number of adherent leukocytes per field of view.

### CFSE

Polyclonal/naïve CD4 T cells, and naive CD8 T cells were purified by Ficoll gradient from peripheral PBMCs of healthy volunteers and stained with CellTrace CFSE (ThermoFischer Scientific) at 1µM as per the manufacturer’s instructions. CFSE stained cells are then cocultured with NECs (HUVECs) or NSCLC-TECs (NSCLC-cocultured HUVECs) for 5 days. Thereafter, cells from the coculture are harvested with TryPLE and stained before being analyzed by flow cytometry.

### Leukocyte and monocyte chemotaxis

PBMCs isolated from the peripheral blood of healthy donors are placed in an 8µm transwell insert and coculture media from monoculture of NECs or NSCLC-TECs is placed in the lower chamber. After 3 hours of incubation, cells that have performed transmigration are collected, centrifuged, and transferred to a conical 96 well plate.

### Spontaneous EC migration assay

After CD31+ sorting, a 96-well plate is coated with gelatin 0.1%, and 5.10^3^ cells per condition are seeded in the wells. The 96-well plate was put in the Incucyte S3 (Sartorius) for live imaging. Phase images were acquired for every experiment. Five images of each well were taken every hour during 24 hours in order to be able to track the cells in a movie. On ImageJ-win 64, using the plugins « Tracking » with « Manual Tracking », the tracking parameters were selected: time interval: 60 min, x/y calibration: 1.24µm, z calibration: 0. 2 cells per image are tracked for 24 hours. Using analysis on Incucyte, the confluence per wall was taken to measure the proliferation of our cells.

### Scratch wound healing assay

Confluent HUVEC monolayers were pretreated for 24 hours with 1µg/ml Mitomycin C to block proliferation. A scratch wound has been made using a 200µl tip. After scratch, cells are washed with PBS and then incubated in supplemented EGM2 in the Incucyte incubator. Photographs were taken every 3 hours for 15 hours and analyzed between T0 and T6. Migration was measured with Fiji and expressed as percentage of wound closure (gap area at T0 minus gap area at T6 in percentage of gap area at T0).

### Quantification and Statistical Analysis

Data represent mean ± SEM between biological replicates. Statistical significance was calculated using non-parametric Wilcoxon test or Kruskal-Wallis depending on the comparison. *p<0.05; **p<0.01; ***<p0.001. Statistical analyses were conducted using GraphPad PRISM.

## Results

### Coculture with NSCLC cells lead to profound transcriptomic EC alterations

To unravel how cancer cells impinge on normal endothelial cells (NECs) behavior we adapted a recent model of 2D coculture ^17^. For this purpose, we selected three NSCLC cell lines (namely A549, H1755, H1975) that encompass the mutational diversity and aggressiveness traits of NSCLC (Figure 1A) and cocultured them directly with a well-established primary cultured model of normal ECs (HUVECs: human umbilical vein ECs) (Figure 1A). Hereafter, HUVEC monoculture is referred to as control normal ECs (NECs), and NSCLC cell-cocultured HUVECs as tumor-like ECs (A549-TECs, H1755-TECs, H1975- TECs). Immunostaining for endothelial markers (vWF and CD31) revealed the spatial organization of HUVEC cocultures as surrounded by tumor cells (Figure 1B). After 48 hours of contact with NSCLC cells, NECs/TECs were enriched using magnetic beads coupled with CD31 antibodies. As expected, flow cytometry analysis of purified cells showed an enrichment for CD31, ranging from 89.9 to 95.8% of CD31+ cells (Figure 1C), while *PECAM1* (the gene encoding for CD31), was not detected in NSCLC by real-time qPCR analysis (Supplementary Figure S1A). Hence, this highlights the purity of the HUVECs purified after coculture with NSCLC cell lines.

**Figure 1.**
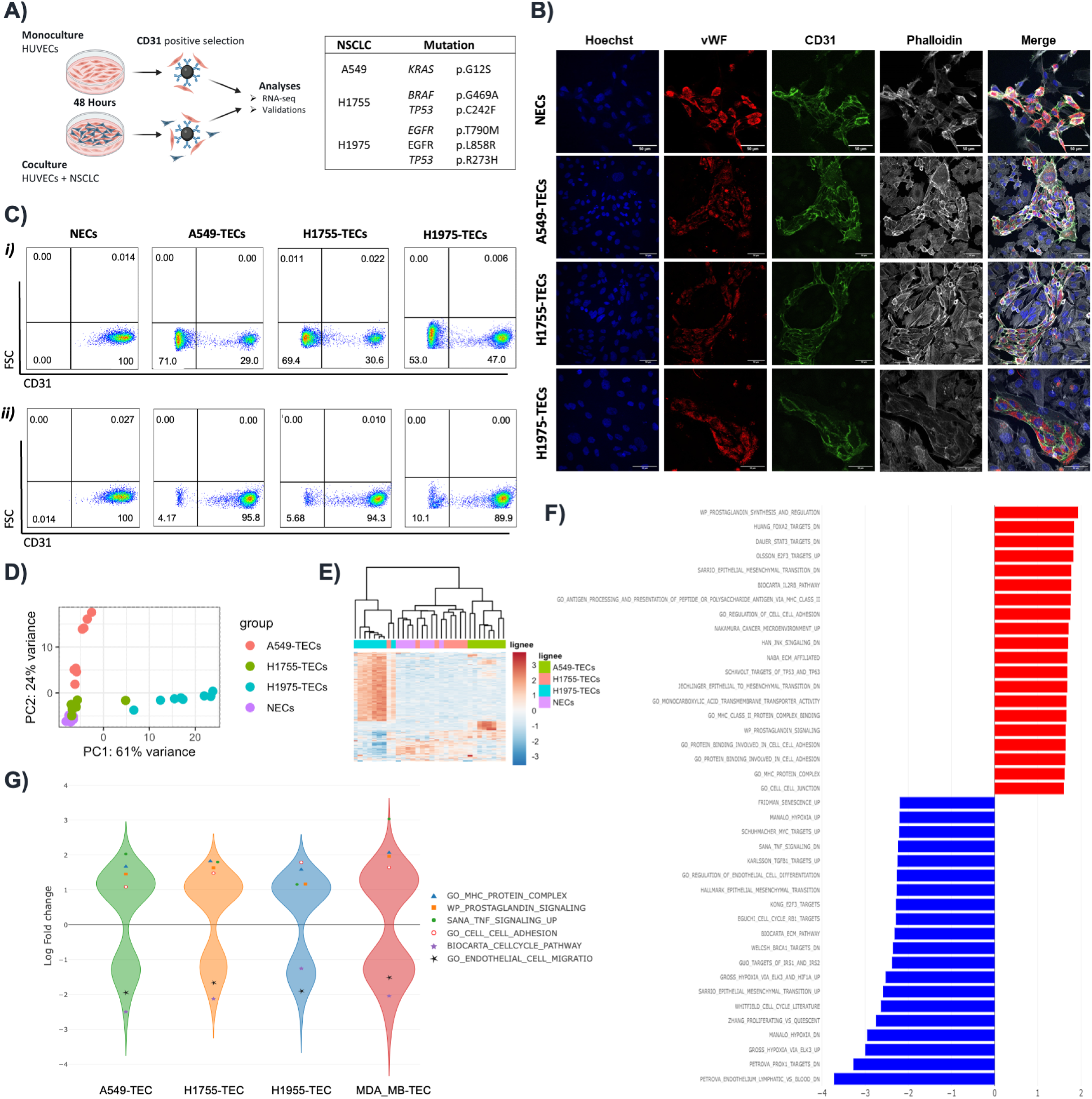
A) Schematic of the 2D in vitro coculture system where human endothelial cells (HUVECs) were cocultured with/without non-small cell lung cancer (NSCLC) cell lines during 48h. Thereafter, ECs were isolated using anti-CD31 antibody-coated magnetic beads and 3’RNA-seq was performed. B) Immunostaining for EC-specific markers (vWF in red and CD31 in green) revealed the spatial organization of HUVECs in cocultures. Scale bar: 50 μm C) Flow cytometric analysis of different cocultured cells stained for CD31 before (i) and after (ii) CD31-sorting to verify population enrichment. D) Principal component analysis considering the top 100 highly-variable genes. Samples segregate into 3 major groups with H1755-TECs being more similar to NECs as compared to the other NSCLC-TECs. n=8 per group. E) Correlation heatmap of the top 350 highly variable genes in NECs and NSCLC- TECs. F) Gene set enrichment analysis in all NSCLC-TECs versus NECs. Pathways enriched in NSCLC-TECs appear in red. p-value p<0.05. G) Violin plot showing results of a meta-analysis between the different NSCLC-TECs and HUVECs cocultured similarly with the breast cancer MDA-MB-231 cell line. Congruent up/downregulated gene sets were observed among the datasets.

We then carried out an unbiased transcriptomics analysis to screen for global changes induced in NECs and TECs. Principal component (PCA) and hierarchical clustering analysis of the highly variable genes revealed that NECs and TECs grouped into 3 distinct clusters (Figures 1D-E). Compared to H1975 and A549 that had a profound impact on the endothelial transcriptomic signature, H1755 minimally affected HUVECs as 4402, 1298 and 137 significantly deregulated genes (adjusted p-value < 0.05) were observed, respectively (Supplementary Fig S1B). Interestingly, H1975- and A549-TECs had 676 commonly deregulated genes, and only 83 genes were in common between the 3 cocultures (Supplementary Fig S1C). Suggesting that difference mediated by NSCLC/EC cocultures could be attributed to distinct genomic alteration profiles. Notably, the impact mediated by NSCLC onto ECs seems to rely on direct cell-cell contacts as indirect coculture (by mean of inserts) did not show any significantly deregulated genes (adjusted p-value < 0.05) and PCA showed that most differences were mediated by the HUVEC donors and not our experimental conditions (Supplementary Fig S1D-E). Gene set enrichment analysis (GSEA) was then performed to identify key upregulated pathways in TECs compared to NECs. Among the 60 significantly upregulated pathways induced upon direct NSCLC cocultures, one third (21 gene sets) suggested a role in EC activation and pro-inflammatory signaling such as the prostaglandin synthesis, TNF-α and complement pathways, IL-22 signaling and leukocyte cell adhesion. On the other hand, gene sets involved in cell cycle, proliferation, EC migration, pseudopodia formation and angiogenesis were downregulated (Figure 1F). When comparing each H1975-, A549 or H1755-TECs to the NECs by GSEA we confirmed unbiased PCA and hierarchical clustering indicating a minimal impact of the H1755-NSCLC cell line on ECs (Figure 1F; Supplementary Fig S1F). Thereafter, to maximize the relevance of our findings, we then performed a transcriptomic meta-analysis with RNA-sequencing dataset of HUVECs cocultured with the MDA-MB-231 breast cancer cell line (hereafter coined as MDA_MB-TECs ^17^). Our meta-analysis revealed congruent deregulated gene signatures between MDA_MB-TECs and NSCLC-TECs as compared to NECs. Indeed, gene sets related to MHC protein complex, prostaglandin signaling, cell-cell (leukocyte) adhesion and TNF signaling were upregulated, while cell proliferation and migration was again downregulated (Figure 1G). Collectively our data indicate that the pro-inflammatory phenotype acquired by ECs upon coculture is probably not a unique feature of NSCLC but could also be generalized to other cancer entities.

### Coculture with NSCLC cells altered EC proliferation and migration

We first investigated the proliferation defect of cocultured HUVECs as suggested by GSEA. Immunostaining of CD31 with the proliferation marker KI67 indeed showed a much lower fraction of double positive CD31+KI67+ cells in our cocultures compared to the monoculture condition (Supplementary Figure S2A). However, when NSCLC-TECs were purified and placed back in culture no significant reduction in cell proliferation was observed, suggesting that the proliferation blockade could be partly mediated by contact inhibition in the NSCLC cocultures (Supplementary Figure S2B). Seemingly, the reduced migration outlined by GSEA was not validated as NECs/TECs displayed similar migration and angiogenesis capacities in spontaneous, scratch wound and tubulogenesis assays (Supplementary Figure S2C-E). Altogether our data suggest that the diminished proliferation and migration phenotype harbored by NSCLC-TECs in cocultures is rapidly lost when ECs are removed from this setting.

### Coculture with NSCLC cells lead to a pro-inflammatory and endothelial activation state

Differential expression analysis and GSEA revealed a marked pro-inflammatory and activation phenotype in all coculture conditions with the expression of multiple cytokines (*CCL26*, *CXCL10, IL16*, *IL18*), and adhesion molecules meant to attract and retain leukocytes (*ICAM1*, *VCAM1, SELE, CD99*) (Figure 2A). We first confirmed by qPCR and flow cytometry the heightened expression of the two major adhesion molecules ICAM-1 and VCAM-1 in NSCLC-TECs (Figure 2B-D). We hypothesized that increased expression of intercellular adhesion molecules by TECs could stimulate the attraction and binding of leukocytes onto the EC surface. Purified NSCLC-TECs were allowed to form a confluent monolayer for 6 and 24 hours, and peripheral blood monocytes (PBMCs) derived from healthy donors were isolated and placed onto NECs/TECs. Interestingly, while there was a trend to increase leukocyte adhesion at 6 hours for the A549-TECs (that expressed the higher level of ICAM-1 and VCAM-1) (Figure 2A-D, Supplementary Figure S3A), this effect was lost after 24 hours of culture. Moreover, at 24 hours the leukocyte adhesion onto H1975- TECs appeared reduced compared to NECs (Figure 2E). As TECs display an altered cytokine profile upon NSCLC coculture, we exposed PBMCs to NSCLC-TEC or NEC secretomes and measured the immune cell chemotaxis with a transwell migration assay. Compared to NEC secretome we observed a trend in increased transmigration of CD3+CD4+ and CD3+CD8+ for all NSCLC-TEC secretomes, with significant results for H1755- and H1975-TECs (Figure 2F). There was no impact on the CD14+ myeloid chemotaxis (Supplementary Figure S3B).

**Figure 2.**
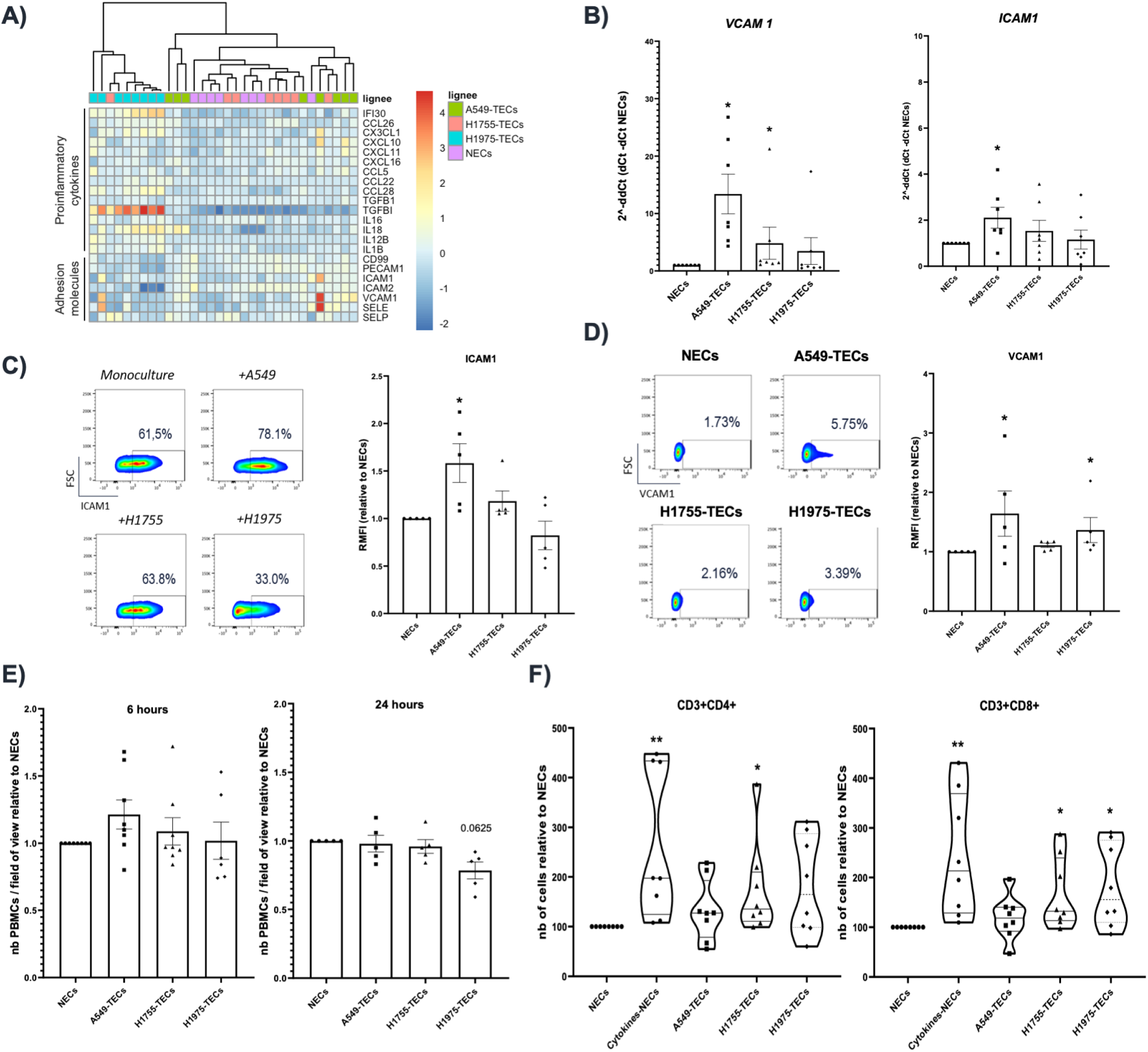
A) Correlation heatmap analysis for genes involved in pro-inflammatory cytokines and adhesion molecules involved in leukocyte recruitment. Expression analysis for VCAM1 and ICAM1 by B) RT-qPCR and C-D) flow cytometry analysis in NECs/NSCLC-TECs. E) NECs and NSCLC-TECs were cultured at confluency for 6h or 24h after coculture and leukocyte adhesion assessed. F) Chemotaxis experiment for CD3+CD4+/CD8+ lymphocytes attracted by the coculture medium from NECs alone or NSCLC-TECs. Data are mean ± SEM, n > 5, *p < 0.05, **p < 0.01, Wilcoxon test compared to NECs.

### NSCLC TECs have limited impact on CD8 T cell activation

Having shown that NSCLC-TECs could promote CD3+ T cell recruitment, we sought to characterize their impact on lymphocyte activation and polarization. Indeed, in several tumor entities including NSCLC, the TME has been shown to induce CD8+ T cell exhaustion, which is the main target of immunotherapies ^21^. After exposing HUVECs or not to NSCLC, NECs/TECs were subsequently cocultured with polyclonal activated CD8+ T cells for 24 hours. In that condition, our flow cytometry analysis did not show significant variation of the late/middle T cell activation marker CD25 in CD3+CD8+ T cells, and only minimal changes in the secretion of TNFα, as measured by ELISA (Supplementary Figures S3C-D).

### NSCLC TECs mediate the polarization of protumor CD4 T cell subsets

The immunosuppression of the TME is also partly imposed by (but not limited to) pro-tumor CD4+ Treg and Th2 polarization ^22^ (Figure 3A). This polarization can be achieved via various cytokines produced by cancer cells (eg. TGFß), but also by direct contact with co-stimulatory molecules expressed on (semi)-professional APC such as ECs ^23^. One of these molecules is OX40L (encoded by *TNFSF4*), which was previously described to be constitutively expressed by ECs ^24^ but never assessed yet in a context of cancer. We first confirmed by qPCR and flow cytometry the reduced expression of OX40L (encoded by *TNFSF4*) in NSCLC-TECs as compared to NECs (Figure 3B-D). Of note, *TNFSF4* expression appeared relatively stable in HUVECs over 72 hours after EC purification (data not shown). OX40- OX40L interaction is described to i) abolish the suppressive activity of FOXP3+ Tregs, ii) prevent the induction of Tregs from effector T-cells, and iii) induce the proliferation of memory and effector T lymphocytes ^25^. OX40/OX40L thus appears as an important immunoregulatory signal, with OX40L expression tightly regulated by several actors including kinases of the PI3K, Src family in mast cell and dendritic cells ^26,27^. To unravel how the OX40L could be regulated in ECs, we performed a kinome assay to screen for 144 serine/threonine kinase (STK) and 196 protein tyrosine kinases (PTK) in TECs compared to NECs. After quality check filtering of the phosphorylated phosphosites and VSN normalization, an upstream kinase analysis algorithm was used to predict differential kinase activity in TECs compared to NECs. Mirroring the results obtained by RNA-sequencing (Figure 1D-E), phylogenetic coral trees showed that A549- and H1975-TECs presented similar modifications in their kinome profile while H1755-TECs have limited effect size compared to NECs (Figure 3E; Supplementary Figure 3E-F). Moreover, the majority of significantly deregulated predicted kinases appeared downregulated in A549- and H1975- TECs, with common kinases belonging to the Src kinase family (LYN, FYN, HCK) (Figure 3E). In immune cells LYN and FYN were previously reported to tightly control OX40-OX40L signaling, with the NFκB pathway that is directly implicated in the OX40L transcriptional expression ^27–29^. As the NFκB heterodimers *NFKB1* and *RELA* (encoding p50 and p65, respectively) were downregulated in H1975-TECs (Supplementary Figure 3G) our data suggest that the downregulation of LYN, FYN kinases and the NFκB signaling could explain the reduced OX40L expression in A549- and H1975-TECs.

**Figure 3.**
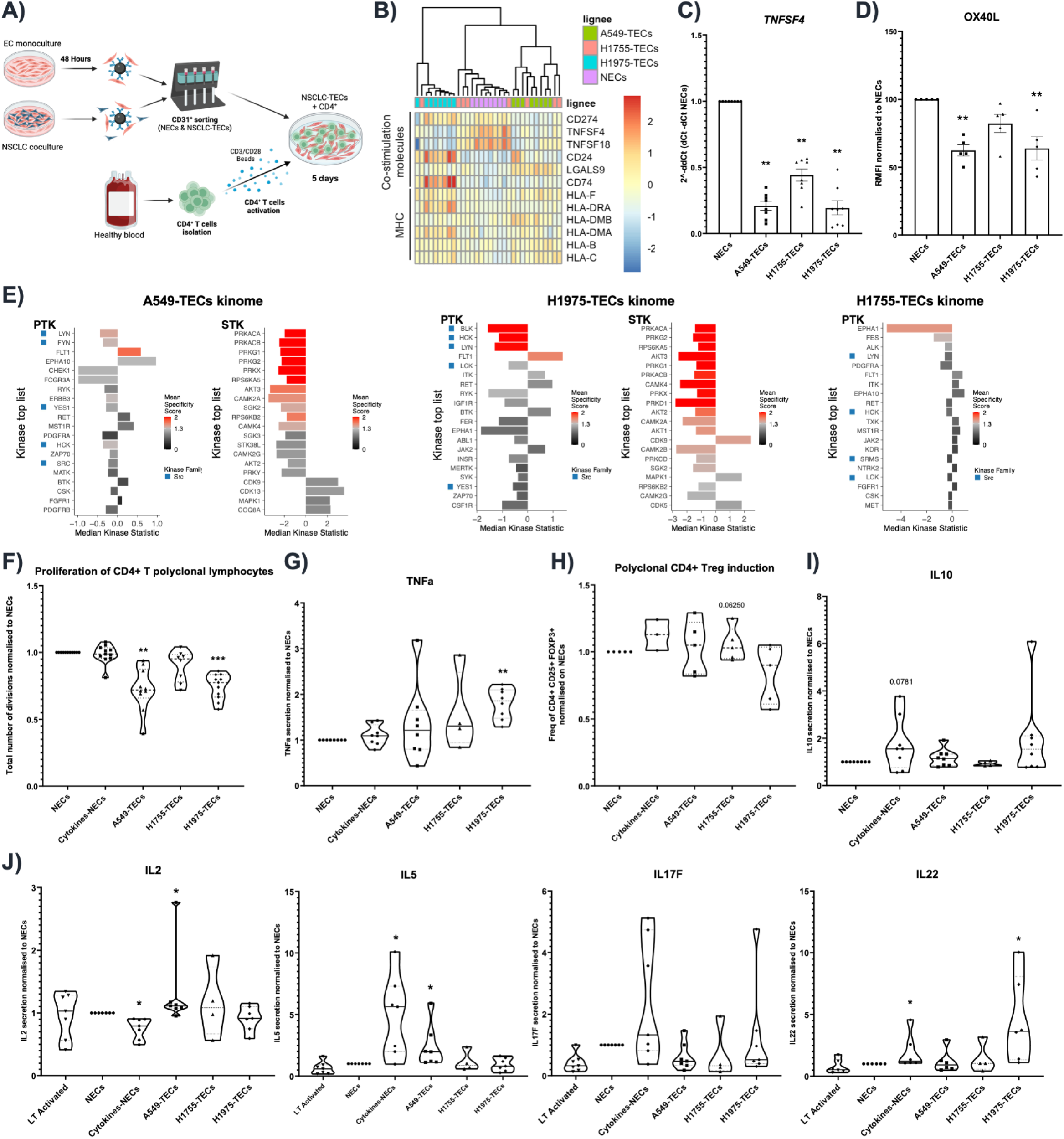
A) Schematic representation of the 5 days coculture between CD4+ T cells and NECs or NSCLC-TECs. B) Correlation heatmap analysis for genes involved in costimulation molecules and major histocompability complexes. Expression analysis for OX40-L (encoded by TNFSF4) by C) RT-qPCR and D) flow cytometry analysis in NEC/NSCLC-TECs. E) Kinome analysis of protein tyrosine (PTK) serine/threonine kinase (STK) differentially regulated in NECs and NSCLC-TECs. A negative median kinase statistic means a downregulated kinase compared to NECs, and a specificity score in red is statistically significant. F) Proliferation of polyclonal CD4+ T cells in cocultures with NECs or NSCLC- TECs and assessed with CFSE by flow cytometry. G) TNFa secretion is dosed by ELISA in the coculture supernatant from polyclonal CD4+ T cells with NECs or NSCLC-TECs. H) Fraction of CD4+CD25+FOXP3+ Treg in the polyclonal CD4+ T cells/HUVEC cocultures. I) Cytokine secretion detected by Legendplex in naïve CD4 T cells/HUVEC cocultures. Data are mean ± SEM, n > 4, *p < 0.05, **p < 0.01, Wilcoxon test compared to NECs.

Following the same strategy as for CD8+ T cells, we scrutinized the impact of NECs/TECs on polyclonal or naïve-activated CD4+ T cells (Figure 3A). When cocultured with NSCLC-TECs, flow cytometry analysis of CFSE-labelled polyclonal CD4+ T cells showed a significantly diminished proliferation with A549- and H1975-TECs (Figure 3F). Conversely, naive CD4+ T cells proliferation was higher when cultured with H1755-TECs compared to NECs (Supplementary Figure S4A). The amount of TNF-α was quantified by ELISA in CD4+ T cell/EC coculture and showed a 1.8-fold change elevation when H1975-TECs were used compared to NECs (Figure 3G). Besides being secreted by activated T cells, TNF-α is described to act on Treg polarization. As such the fraction of CD4+CD25+FOXP3+ Treg was higher with A549- and H1975-TECs for polyclonal CD4+ T cells but not affected in naive CD4+ T cells (Figure 3H; Supplementary Figure S4B). Seemingly, there was a tendency for increased IL-10 secretion in cocultures with polyclonal CD4+ T cells and A549- or H1975- TECs (IL-10 being a cytokine secreted by Treg) (Figure 3I). Using the LegendPlex technology we then performed a thorough investigation of the cytokines produced by CD4+ Treg and T helpers in the cocultures of naive CD4+ T cells with ECs. Notably, in monoculture upon TNFα/IL-1ß stimulation, naïve CD4+ T cells seem to polarize in Th2 (IL5+, IL13+, IL2-) and Th17 (IFNG+, IL22+) (Figure 3J; Supplementary Figure S4C). Our coculture results indicate that A549-TECs could induce the secretion of IL-2 and IL-5, secreted by Th1 and Th2 respectively (Figure 3J). On the other hand, H1975-TECs only fostered IL-22 secretion, with low IL-17 levels suggesting a Th22 polarization (Figure 3J) ^30,31^. Notably, IL-4, IL-13 and IFNγ showed minimal variation (Supplementary Figure S4C), and IL-9, IL-10 and IL-17A were undetectable. A combination of various polarization factors is described to mediate a specific CD4+ subtype. For instance, IL-6 and TNF-α (without TGFß) that stimulate the differentiation of Th22 ^32^, are upregulated in H1975-TECs (Figure 3G; Supplementary Figure S4D-E). Hence, collectively our data point toward a Th22-polarizing effect of H1975-TECs, and a mixed Th1/Th2 signature with A549-TECs.

### NSCLC-TECs mediate the polarization of anti-tumor M2-like macrophages

The NSCLC immune TME is mostly composed of myeloid cells including monocyte-derived macrophages, that have pivotal functions in tumor immunity ^33^. To unravel the impact of ECs during this process we cocultured NSCLC-TECs and NECs with monocytes isolated from healthy peripheral blood (Figure 4A). After 5 days of culture, confocal microscopy analysis of monocyte/NEC cocultures revealed that CD45+CD163+ monocytes appear enriched in areas of close contact with NECs (Figure 4B). We then further analyzed by flow cytometry monocyte/NEC or TEC cocultures, and particularly the CD31-CD45+ cell fraction for accepted pro-tumor M2-like (CD163, CD16) and anti-tumor M1-like macrophage markers (HLA-DR). Cocultures between NECs/TECs and monocytes had limited impact on the monocyte polarization. Nonetheless, we observed a propensity toward an increased expression of CD163 and HLA-DR with A549- and H1975-TECs, respectively (Figure 4C; Supplementary Figure 4F).

**Figure 4:**
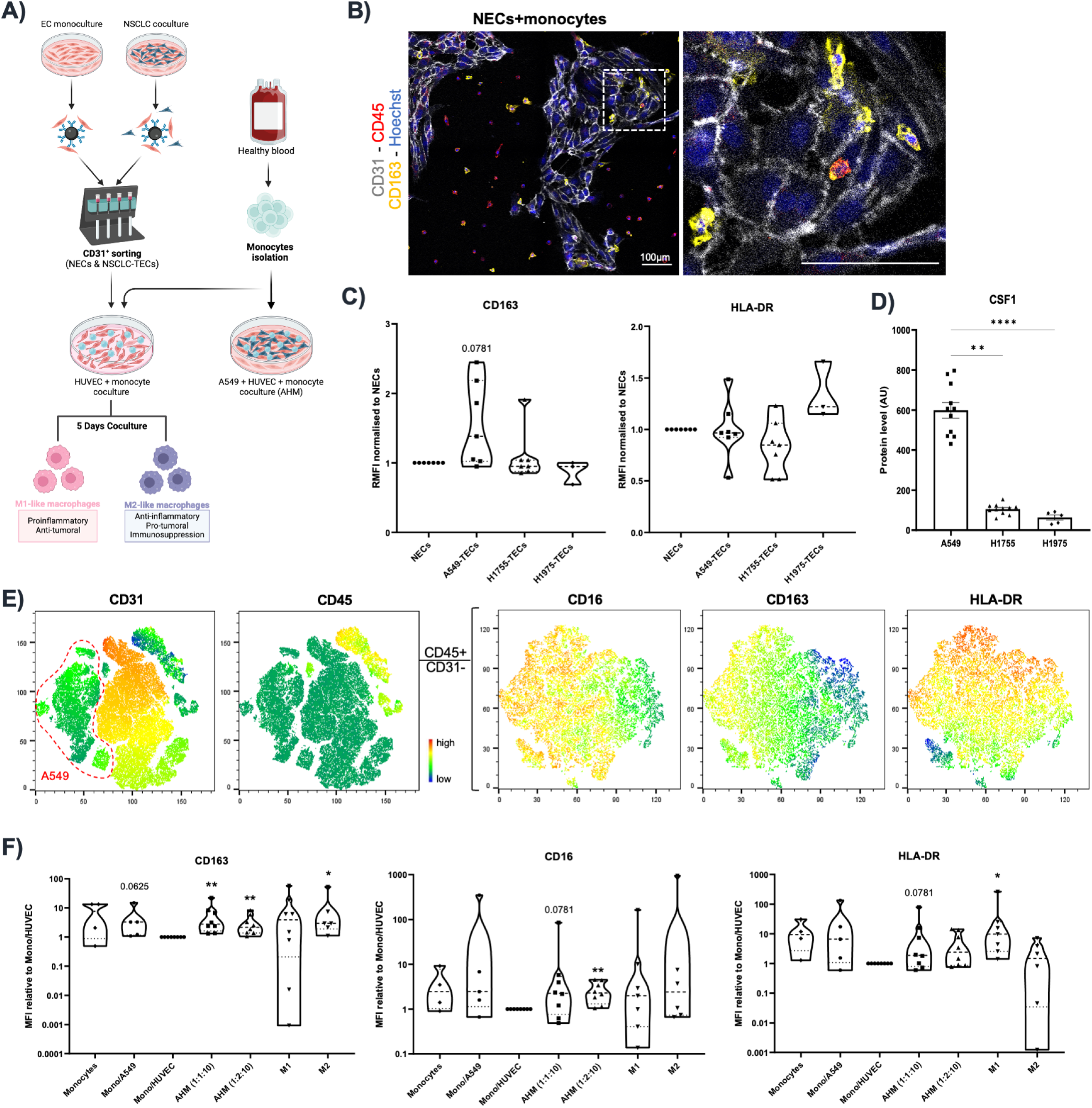
A) Schematic representation of the 5-day coculture between PBMC-derived monocytes and NECs or NSCLC-TECs. B) Confocal analysis of the coculture between healthy HUVECs (NECs) and monocytes, and stained for the endothelial marker CD31, the leukocyte marker CD45, and the M2-like macrophage marker CD163. Scale bar: 100μm. C) Flow cytometry analysis of several polarization markers after the coculture between monocytes and NECs or NSCLC-TECs. HLA-DR is a M1-like marker. D) Mass spectrometry proteomic analysis of the secretome of NSCLC cell lines for CSF1. E) tSNE representations and F) mean of fluorescence (MFI) of flow cytometry of the AHM triculture with A549, HUVEC and monocytes. The right panels are gated on the CD45+CD31- population to study macrophage polarization for M2-like (CD16 and CD163) and M1-like (HLA-DR) markers. Data are mean ± SEM, n > 4, *p < 0.05, **p < 0.01, Wilcoxon test compared to Mono/HUVEC.

Previous studies from our lab showed that thoracic cancer cell lines could mediate the M2-like polarization of monocytes in 2D and 3D cocultures, involving tumor-secreted cues including the M-CSF/CSF1 ^34–36^. Interestingly, mass spectrometry analysis demonstrated a higher expression of CSF1 in A549 secretome compared to those of H1755 and H1975 (Figure 4D). Hence, to combine the impact of NSCLC on monocyte polarization, we directly tri-cultured in 2D the A549 NSCLC cell line with ECs (HUVECs) and monocytes for 5 days (Figure 4A). From now on, to simplify reading we will dub the 2D A549/HUVECs/monocyte triculture as AHM. We then screened by flow cytometry the AHM triculture by gating on the CD45+ immune population that was well defined on TSNE representations (Figure 4E). First, when monocytes were cultured with A549 we reproduced the heightened CD163 expression (and CD16, although not significant) as seen with other thoracic cancer cell lines ^34,36^. Second, the addition of ECs in the AHM triculture did not alter the increased expression of CD163, even when using a higher ratio of ECs in the triculture, but the levels of CD16 were boosted. Third, the addition of ECs was however able to induce the expression of the M1-like marker HLA-DR, but a greater EC ratio did not impose more variation in this marker (Figure 4F). As such, our data suggest that ECs could favor the emergence of a pro-tumor M2-like phenotype whilst minimally affecting the M1-like phenotype.

### A multicellular tumor spheroid (MCTS) 3D model to mimic the lung TME

In the past decade, new culture systems have emerged, accounting for the study of 3D environment and that appear more similar to *in vivo* setting as compared to classical 2D monocultures ^37,38^. Hence, in order to complexify our coculture systems, we modeled the TME with 3D MCTS. MCTS were obtained by spontaneous cell aggregation over 5 days with a defined ratio (2:1:1:1) of A549, HUVECs, PBMC-derived monocytes isolated from the peripheral blood of healthy donors, and primary healthy lung fibroblasts (Figure 5A). As previously published by others ^39^, we showed that the use of fibroblast was necessary for MCTS condensation and formation, probably by secreting important components of the extracellular matrix (ECM) (data not shown). Confocal microscopy imaging revealed an elaborated cellular organization with FAP+ fibroblasts forming a network in the center of MCTS, embedding clusters of CD31+ ECs and CD45+ monocytes where those cells appeared to form privileged contacts (Figure 5B). To further unravel this complexity, MCTS were dissociated into a highly viable (>85%) single cell suspension, subsequently analyzed by single cell RNA-sequencing (scRNA-seq). After quality filtering and integration (Supplementary Figure 5), we obtained 52.206 high-quality cells that clustered into 6 distinct clusters, annotated on the basis of their top-50 marker genes (Figure 5C-D; Supplementary Figures 6A-B). Cancer cells were identified by the expression of epithelial cell markers and genetic alterations as outlined by inferring somatic large-scale chromosomal copy number alterations with inferCNV (Supplementary Figure 6C). This step was necessary to formally distinguish cancer cells and fibroblasts that share similar features. The 31.878 cancer cells fall into 5 subclusters with specific transcriptomic signatures such as a proliferative subcluster, cholesterol and fatty acid homeostasis, and EMT/inflammatory signaling (Supplementary Figure 7). Using transcription Single-Cell Regulatory Network Inference and Clustering (SCENIC), we could detect enriched activity of CEBP and c-JUN-driven regulons and transcription factor expression, congruent with the proliferative capacities of cancer cells (Supplementary Figure 8).

**Figure 5:**
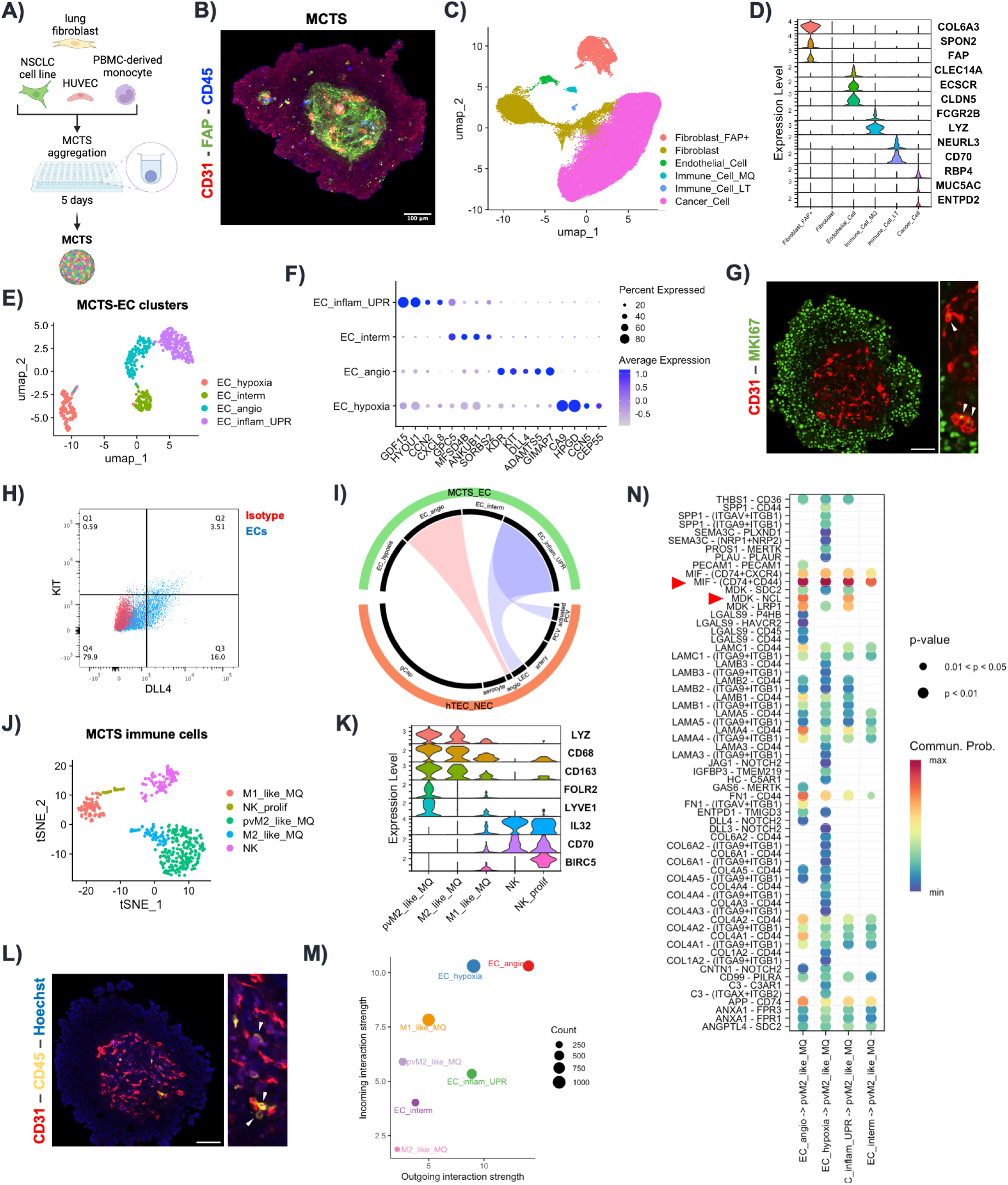
A) Schematic representation of the multicellular tumor spheroid (MCTS) formation during 5 days. B) Confocal imaging of MCTS for the cancer-associated fibroblast (CAF) marker FAP, the leukocyte marker CD45 and the endothelial CD31. Scale bar, 100μm. C) uMAP representation of each cell type composing the MCTS and detected after scRNA-sequencing (n=4). D) Violin plot of key marker genes for each cell type in the MCTS. Subclustering analysis of the MCTS-ECs visualized by E) uMAP with F) dotplot depicting key marker genes. We annotated inflammatory/UPR (unfolded protein response), intermediate, angiogenic and hypoxia (proliferative) EC subsets. G) Confocal imaging of MCTS for CD31 and the proliferative marker MKI67. Arrow heads indicate proliferative CD31 cells. Scale bar, 100μm. H) Cytometry analysis of MCTS for angiogenic markers KIT and DLL4, as compared to control isotypes. I) Circleplot showing the similarities between MCTS-EC subsets and ECs freshly isolated from NSCLC human biopsies. Subclustering analysis of the MCTS immune compartment visualized by J) tSNE with K) violin plot depicting key marker genes. L) Confocal imaging of MCTS for CD31 and the immune marker CD45. Arrow heads indicate close interactions between CD45 monocytes and CD31 cells. Scale bar, 100μm. M) Ligand-receptor predicted the interactions between MCTS-EC and the myeloid compartment present within MCTS. N) Heatmap of the top predicted pathways involved between the perivascular M2-like macrophage (pvM2-like) and the MCTS-EC clusters. The most significant pathway relates to the MIF (macrophage migration inhibitory factor) and MK (midkine).

Although healthy fibroblasts were used, among the 19.389 fibroblasts 18% expressed the cancer-associated fibroblast (CAF) marker FAP and were shared across all samples, while normal fibroblasts displayed heterogeneity between donors ^40^ (Supplementary Figures 9A-B). Heterogeneous phenotypes were identified thanks to gene signatures from breast cancer scRNA-seq data ^41^ including immunosuppressive S1_ECM_myCAF and S1_TGFb_myCAF with a low relative fraction of immunoactivatory S1-detox-iCAF (Supplementary Figure 9C).

With respect to ECs isolated from MCTS (coined MCTS-ECs), the 505 cells obtained post filtering segregated into 4 distinct subclusters (Figure 5E). Probably linked to the hypoxic nature of the central core of MCTS, we found a first hypoxic EC cluster characterized by the hypoxic marker *CA9* and the prostaglandin-degrading *HPGD* that was closely associated with capillary aerocytes in human lungs ^42^. Probably linked to the hypoxic response, this MCTS-EC cluster also expressed established proliferation markers (*TOP2A, MKI67, BIRC5*, *CEP55*) and pathways related to ECM remodeling. A second MCTS-EC subcluster, coined angiogenic MCTS-ECs expressed genes involved in angiogenesis (*KDR*, *FLT1*, *TIE1, KIT*), with tip-like markers (*INSR*, *ANGPT2*, *ANGPTL2*) and ECM remodeling (*COL4A1, COL4A2, COL18A1*), as described previously ^9^. Of note, the proliferative MCTS-EC subcluster also express some of angiogenic markers, probably suggesting a stalk-like phenotype. A third MCTS-EC subcluster had a mixed intermediary phenotype with a poorly defined signature. It was thus coined as intermediary (Figure 5F). Using confocal imaging and flow cytometry we have validated some of the main markers from the proliferative and angiogenic MCTS-EC clusters (Figure 5G-H). Finally, the fourth MCTS-EC subcluster displayed a marked pro-inflammatory signature with genes encoding for cytokines (*CXCL1*, *CXCL2*, *CXCL8*), NFKB pathway (*CCN2*), leukocyte adhesion molecule (*ICAM1*), endothelial-to-mesenchymal transition (*SERPINE1*), and endoplasmic reticulum stress-response (*HSP90B1*/GP96, *HSPA5*/BIP, *PDIA4*, *SELENOK/S*/*M*, *DNAJC*, *SERP1*, *HERP, HYOU1*) (Figure 5F). GSEA further highlighted the implication of the UPR, autophagy and mTORC1 signaling in the inflammatory/UPR MCTS-ECs, as recently characterized in mouse TECs ^11,12^. We then sought to unravel the relevance of those MCTS-EC phenotypes by determining their similarities across models. While the proliferative and angiogenic MCTS-ECs were found in scRNA-seq datasets of 2D monoculture of normal HUVECs ^43^, the inflammatory/UPR MCTS- EC subcluster was absent in early 2D culture of HUVECs (Supplementary Figure 10A). Conversely, the inflammatory/UPR MCTS-EC subcluster was expressed in freshly-isolated ECs from NSCLC human biopsies and appeared similar to ECs with immunomodulatory functions such as lymphatic ECs and activated postcapillary venule (Figure 5I). Similarly, the inflammatory/UPR MCTS-EC subcluster was found similar to a cluster expressing a mesenchymal/EndMT signature found in 2D culture of TECs from NSCLC biopsies ^9^ (Supplementary Figure 10B).

We then further phenotyped the immune cells of MCTS clusters, originating from PBMC-derived circulating monocytes. Subclustering analysis revealed a formidable heterogeneity with 434 cells that cluster into 5 subclusters (Figure 5J). Notably, probably due to contamination inherent to the monocyte isolation, 2 clusters of natural killer (NK) cells were identified, expressing *CD96*, *IL32, CD70* and *RELB* (Figure 5K). They were labeled as NK, and NK proliferative by interrogating reference-based atlas ^44^. Those lymphoid clusters will not be further analyzed in this study. Emphasizing on macrophage diversity during differentiation and activation processes in the TME, 3 heterogeneous cluster of myeloid cells were identified on the basis of canonical markers (*LYZ*, *CD86*, *CD14*, *CD68*) (Figure 5K). Although a continuum exists within these subsets, we detected i) a perivascular M2-like macrophage characterized by *CD163* and the accepted markers *LYVE1* and *FOLR2* ^45^; ii) a M2-like phenotype with high expression of M2-like markers but no expression of *LYVE1* and *FOLR2* ; and iii) a population of M1-like macrophage expressing low levels of the aforementioned markers (Figure 5K). Interestingly, the M2-like fraction represents 77% of the myeloid compartment, mirroring our previous observations in NSCLC and mesothelioma MCTS that most of the monocyte-derived macrophages adopt an M2-like phenotype ^35,36^. Given that CD45+ were found in the close vicinity of ECs within MCTS (Figures 5L), we predicted ligand-receptor interactions between MCTS-immune and ECs with CellChat to identify therapeutically-relevant targets. Notably, the angiogenic MCTS-EC subcluster was predicted to perform most interactions (both incoming and outgoing), with the highest weight (Figures 5M; Supplementary Figure 10C). Interestingly, the macrophage migration inhibitory factor (MIF) and the midkine (MK) pathways were the pathway showing the most significant interactions in the perivascular M2-like cluster (Figure 5N). MIF was identified as a ligand of both CXCR2 and CXCR4, that are major regulators of inflammatory cell recruitment ^46^ (Figure 5N; Supplementary Figure 10D-F). Moreover, another receptor of MIF is CD74, involved in TNF-α–dependent lung inflammation ^47^, which was highly expressed in M2-macrophages subsets (Supplementary Figure 10F). Finally, the MK signaling was recently identified in the immunoregulatory of the endometrial carcinoma ^48^. Our data predict that the inflammatory/UPR and angiogenic MCTS-EC clusters are the major endothelial producers of MK, which could act on M1- and perivascular M2-like macrophages (Supplementary Figure 10D-F). Altogether, this data supports the evidence that MCTS-ECs could regulate tumor macrophage polarization via various pathways, and that MCTS appear as a relevant model to mimic some aspects of cell-cell communications from the TME.

## Discussion

The aim of our study was to model the TME and analyze the impact of ECs on various immune populations, in order to better dissect their immunoregulation function. We showed that coculturing ECs with various NSCLC cell lines induced profound alterations at the transcriptomic, proteomic and kinomic levels with the induction of pro-inflammatory pathways, reflecting their activation state. Interestingly, the pro-inflammatory phenotype appears mainly driven by direct cell contact as indirect coculture with inserts had minimal effect. Besides, this feature does not appear restricted to the NSCLC entity, as our meta-analysis revealed a similar transcriptomic signature in breast cancer cell cocultured-ECs. Despite a limited impact of NSCLC-TECs on CD8 T lymphocyte proliferation and activation, we showed that CD4 T cells were polarized into different subsets (Treg, Th22 and Th1/Th2) when cocultured with NSCLC-TECs. We also discovered that NECs and NSCLC-TECs could enhance the expression of M2-like markers. For the first time, we brought to light that OX40L could play a role in the immunoregulatory function of TECs, where it appears downregulated as compared to NECs.

In the seek to improve our 2D coculture system, we scrutinized at the single cell resolution, 3D MCTS encompassing tumor cells, ECs, fibroblasts and monocyte-derived macrophages. We showed that cell heterogeneity was much more complex than 2D cultures, with several cell phenotypes identified. Particularly, 3 subclusters of MCTS-ECs were distinguished, among which the inflammatory subcluster that was absent from standard 2D culture but present in freshly-isolated ECs from NSCLC biopsies. Finally, we uncovered various macrophage polarization states within NSCLC-MCTS, with a perivascular M2-like subset that has particular therapeutic importance with regard to its interaction with the vascular compartment.

Our results on OX40L raise several outstanding questions and promising future perspectives. OX40L/gp34 belongs to the TNF family and is identified as the ligand for the costimulatory receptor OX40. Importantly, the use of an anti-OX40 agonist antibody has been shown to enhance T lymphocyte-mediated antitumor immunity and inhibit Treg induction in several cancer models, thereby promoting tumor regression and ameliorating patient survival ^49,50^. Moreover, OX40L has been identified to be expressed in ECs (HUVECs) where it can act as a co-stimulation signal for T cell recruitment and proliferation ^24,51,52^. Here we show that OX40L expression was blunted in A549- or H1975-TECs, and their coculture with polyclonal CD4 T cell inhibits their proliferation following activation, while increasing Treg proportion. We hypothesize that inhibition of CD4 T cell proliferation results from the failure of the proliferative support signal normally provided by OX40/OX40L at the time of T cell activation. Indeed, this hypothesis is reinforced by the fact that no difference in proliferation was observed in naive T cells, which only express OX40 very late after activation ^53^. An important interrogation holds in deciphering the signaling leading to reduced OX40L expression in some NSCLC-TECs compared to NECs. In various immune cell types, the OX40/OX40L signaling appears regulated by the NFκB pathway ^27–29^. Moreover, in HUVECs the OX40 binding results in c-Jun and c-Fos induction (10477563) that closely interact with the NFκB p65 subunit, thereby mutually activating their binding to DNA motifs ^54^. Here, we show that the expression of the Lyn and Fyn kinases (involved in NFκB pathway regulation ^29^) is reduced in A549- or H1975-TECs, and might be implicated in the OX40L diminution but this hypothesis needs be further tested. Noteworthy, given the role of the OX40/OX40L interaction in the establishment of an effector memory response ^25^, it will be relevant to examine effector (CD45RA-CCR7-) and central (CD45RA-CCR7+) memory T cell markers after activation upon coculture with NSCLC-TECs.

The characterization of IL-22 Th22 cells is relatively recent and initially defined as cells specializing in tissue repair processes. Intra-tumor IL22-producing Th22 cells, which proportion is increased in tumor tissues, is associated with poor prognosis in several cancer types ^55–57^. Here we show that naive CD4 T cells cocultured with H1975-TECs fostered IL-22 secretion, while producing low IL-17 levels and thus suggesting a Th22 polarization ^30,31^. However, the effect of A549-TECs appears less clear compared to the control condition stimulated with TNFα/IL-1ß where naïve CD4+ T cells polarize in Th2. Indeed, although IL-5 secretion was induced upon coculture with A549-TECs, there was only a tendency toward heightened IL-13, and the IL-2 (a Th1 marker) was also induced. We hypothesize that this incomplete Th2 signature might relate to the coculture duration and would require later end point analysis. A mass spectrometry proteomic characterization of NSCLC-TECs secretome may also identify polarization mediators eventually involved in this mixed Th2 phenotype. Placing our results back in the context of the TME, we thus hypothesize that TECs may contribute to the polarization of Th2 and Th22 cells, hence participating in tumor immunosuppression. Following on from this project, it would be interesting to deepen our understanding of the underlying mechanisms involved in the interaction between ECs and Th2/Th22 in normal and tumor contexts.

In NSCLC, as it is the case for many other cancer entities, TAMs represent the major immune cell type, accounting for more than one third of tumor-infiltrating immune cells ^33^. Macrophages display high intrinsic plasticity and adaptability based on epigenetic regulation and various cues emanating from the TME. Beyond the canonical M1/M2 macrophage phenotype dichotomy, scRN-seq emphasized macrophage diversity in cancers (30979687). As such, macrophages exist along a phenotypic continuum and are not easily characterized by individual markers ^58,59^. As shown by our lab and others, monocytes differentiate toward M2-like macrophages when cultured in 3D with tumor cells ^34–36^. Here, we show by scRNA-seq a much more in-depth resolution of this macrophage continuum with several intermediate phenotypes. In 2D, when cultured with ECs, macrophages seem to express high levels of CD163 when located nearby ECs. Seemingly, in MCTS CD45+ monocytes appeared to form privileged contact with MCTS-ECs. These observations are consistent with the results from Njock et al. which indicate that treatment with extracellular vesicles produced by HUVECs during coculture with MDA-MB-231 induced polarization towards an M2 phenotype ^17^. Finally, our MCTS data identify the presence of LYVE1+ FOLR2+ perivascular M2-like macrophages described to have therapeutic relevance by influencing treatment resistance, angiogenesis, EC-leukocyte interactions and patient survival ^60–63^. Although protein validations will be required to firmly establish their true nature it will be interesting to determine whether the presence of ECs alone suffices to induce M2-like perivascular traits or whether it requires other signals from the TME.

We acknowledge limitations of our study. First, we restrained our 2D and 3D coculture systems to a single EC type with the HUVEC, but ECs from other vascular beds may show different responses to the contact with tumor cells. For instance, we and others ^9^ showed that normal aerocyte and general capillaries appear downregulated in NSCLC tumors. As such tumor cells or the TME may have a particular detrimental effect on this EC type. Second, in our study we determined the effect of NECs/TECs on the tumor immune compartment, as well as how tumor cells could influence ECs during this process. In fact, cellular communication within the TME is not unidirectional, and lymphocytes, macrophages or fibroblasts also have impacts on ECs ^64,65^. Third, several targets and cell-cell predictions will require to be thoroughly validated and visualized using high dimension in situ imaging. Fourth, the 2D cocultures and MCTS cannot be seen as ideal models to mimic the TME. Rather they are means to study specific cell type/phenotype in which this culture model appears similar to the *in situ* condition. Nevertheless, adding flow to the MCTS with microfluidic device could certainly increase EC heterogeneity with artery/vein specification.

Notwithstanding these limitations, we believe our various culture models allow to better delineate the precise function of TECs within the TME, and how they could modulate tumor immunity. In the future, the MCTS could represent an added-value as a platform for drug screening and to dissect in-depth aspects of immune related EC functions.

## Acknowledgments

We thank Eric Letouze, Marie Denoulet and Jennifer Derrien for valuable discussion on the bioinformatic analysis.

## Supplementary figure legends

**Supplementary Figure 1:**
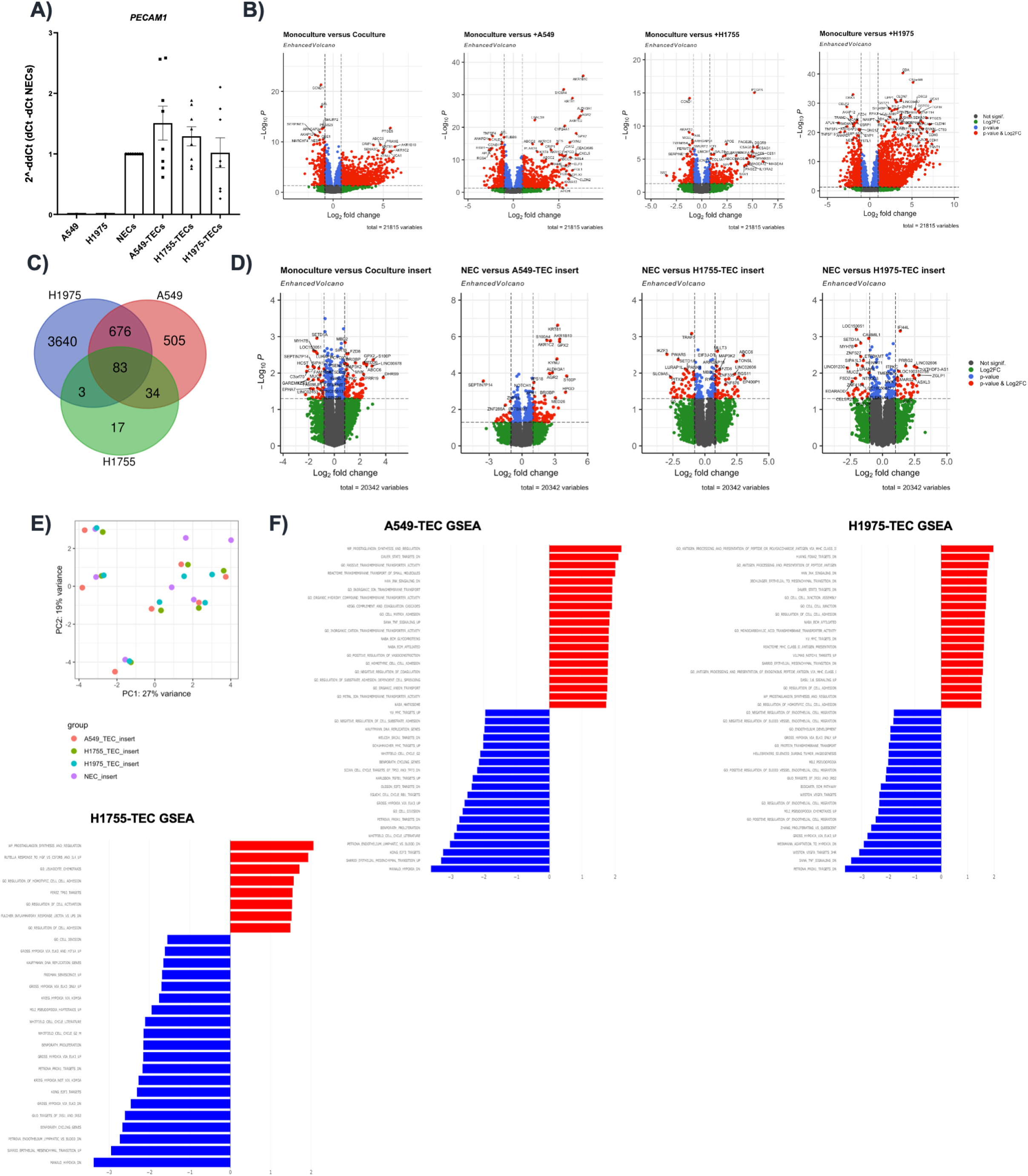
A) RT-qPCR analysis of NSCLC cell lines and NSCLC-TECs after enrichment for the endothelial marker PECAM1. B) Volcano plots depicting differential analysis between each individual NSCLC direct coculture relative to the HUIVEC monoculture (adjusted p-value p<0.05). C) Venn diagrams indicating congruent deregulated genes between each coculture versus monoculture (adjusted p-value p<0.05). D) Volcano plots depicting differential analysis for the indirect cocultures system. E) PCA showing the distribution of each sample in the indirect coculture assay. Clustering appears mediated by the different HUVEC donors and not by experimental conditions. F) Gene set enrichment analysis in each individual NSCLC-TECs versus NECs. Pathways enriched in each NSCLC- TECs appear in red. p-value p<0.05.

**Supplementary Figure 2:**
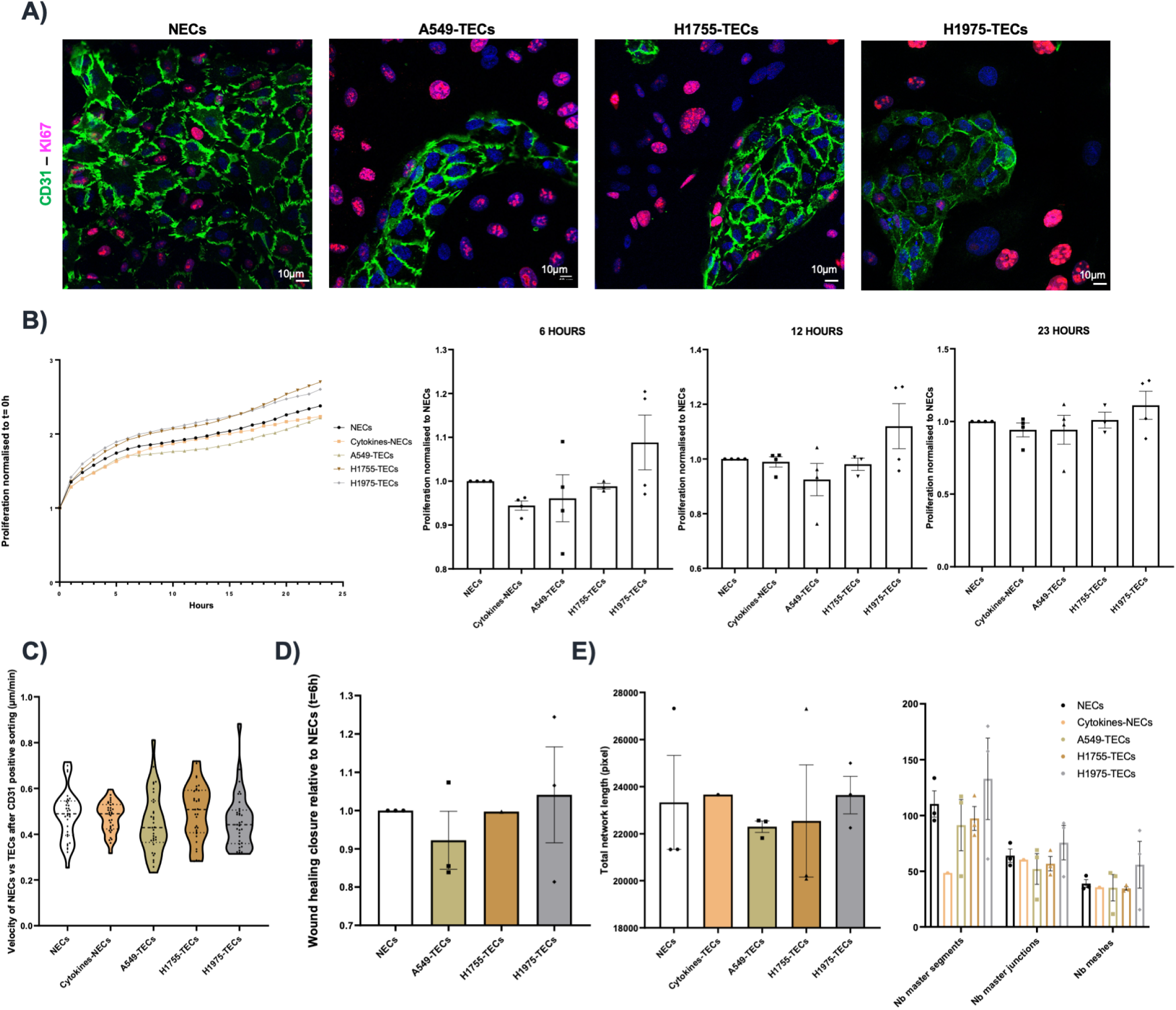
A) Confocal microscopy of NECs and NSCLC-TECs cocultures, stained for the endothelial CD31 and the proliferation marker KI67. NECs and NSCLC-TECs were enriched with CD31 after cocultures and screened for various functions upon culture. B) Proliferation, normalised to t = 0h in function of time and at different times (t = 6h, 12h and 23h) normalized to monoculture. C) Velocity, in μm/min and normalized to monoculture. D) Scratch wound closure, and E) 2D tubulogenesis assay. Data are mean ± SEM, n > 3, Wilcoxon test compared to NECs.

**Supplementary Figure 3:**
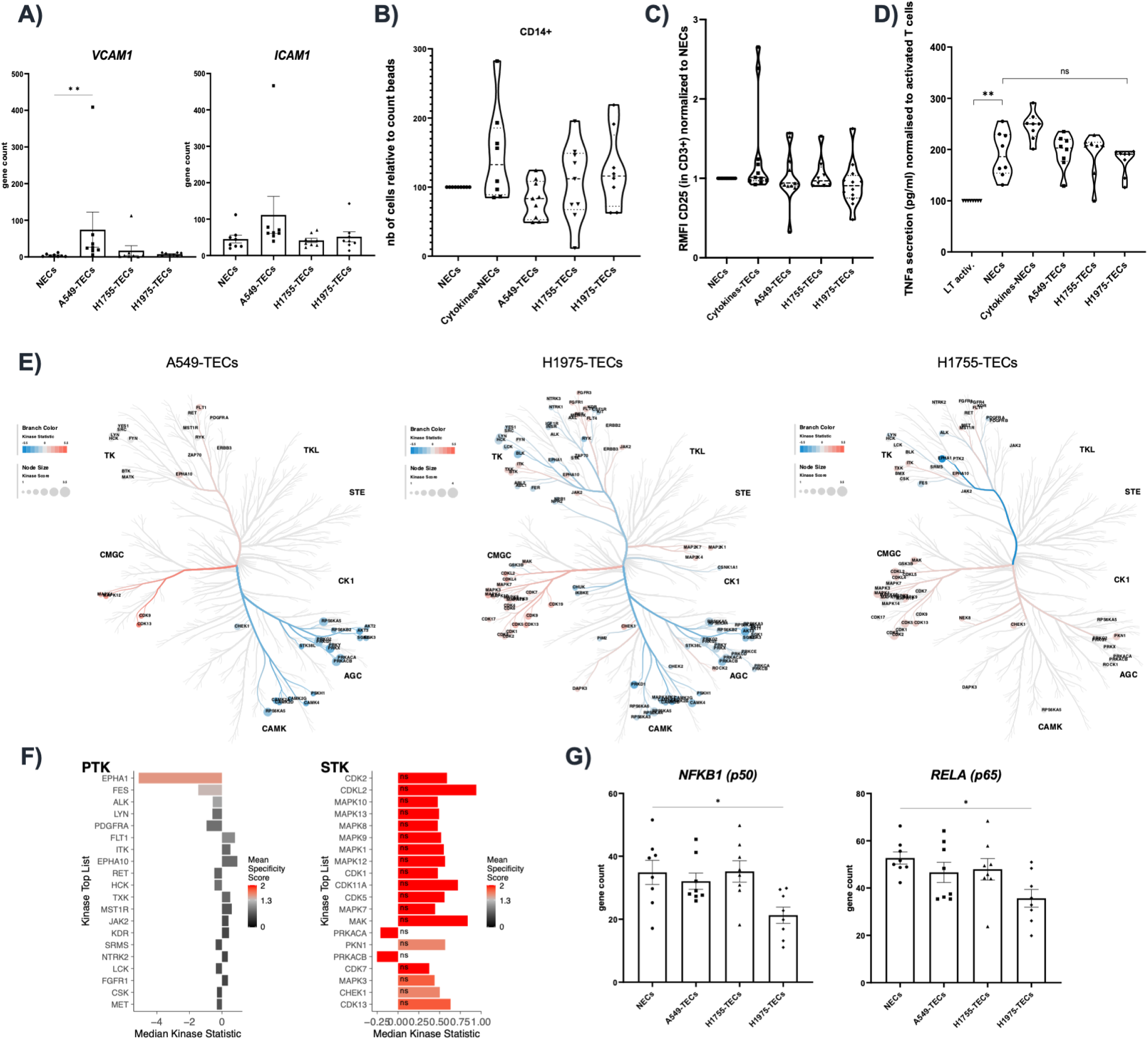
A) Gene count expression from the 3’RNA-seq for VCAM1 and ICAM1 in NECs and NSCLC-TECs. B) Chemotaxis experiment for CD3+CD4+/CD14+ monocyte attracted by the coculture medium from NECs alone or NSCLC+TECs. C) Flow cytometry analysis for CD25 and D) TNFα ELISA secretion in polyclonal activated CD8+ T cells cocultured with NEC/NSCLC-TEC. E-F) Kinome analysis of protein tyrosine (PTK) serine/threonine kinase (STK) differentially regulated in analysis in NECs and NSCLC-TECs. E) Coral trees of the kinase differentially regulated in NSCLC-TECs compared to NECs. Kinase statistic represent the expression of the kinase compared to control (red is upregulated, blue downregulated). The diameter of the dots is proportional to its significativity. F) Bar plots of the deregulated kinases in H1755-TECs. Note that all STK are not statistically significant. G) Gene count expression from the 3’RNA-seq for NFKB1 and RELA in NECs and NSCLC-TECs. Data are mean ± SEM, n > 8, *p < 0.05, **p < 0.01, Kruskal-Wallis test, and Wilcoxon test compared to NECs.

**Supplementary Figure 4:**
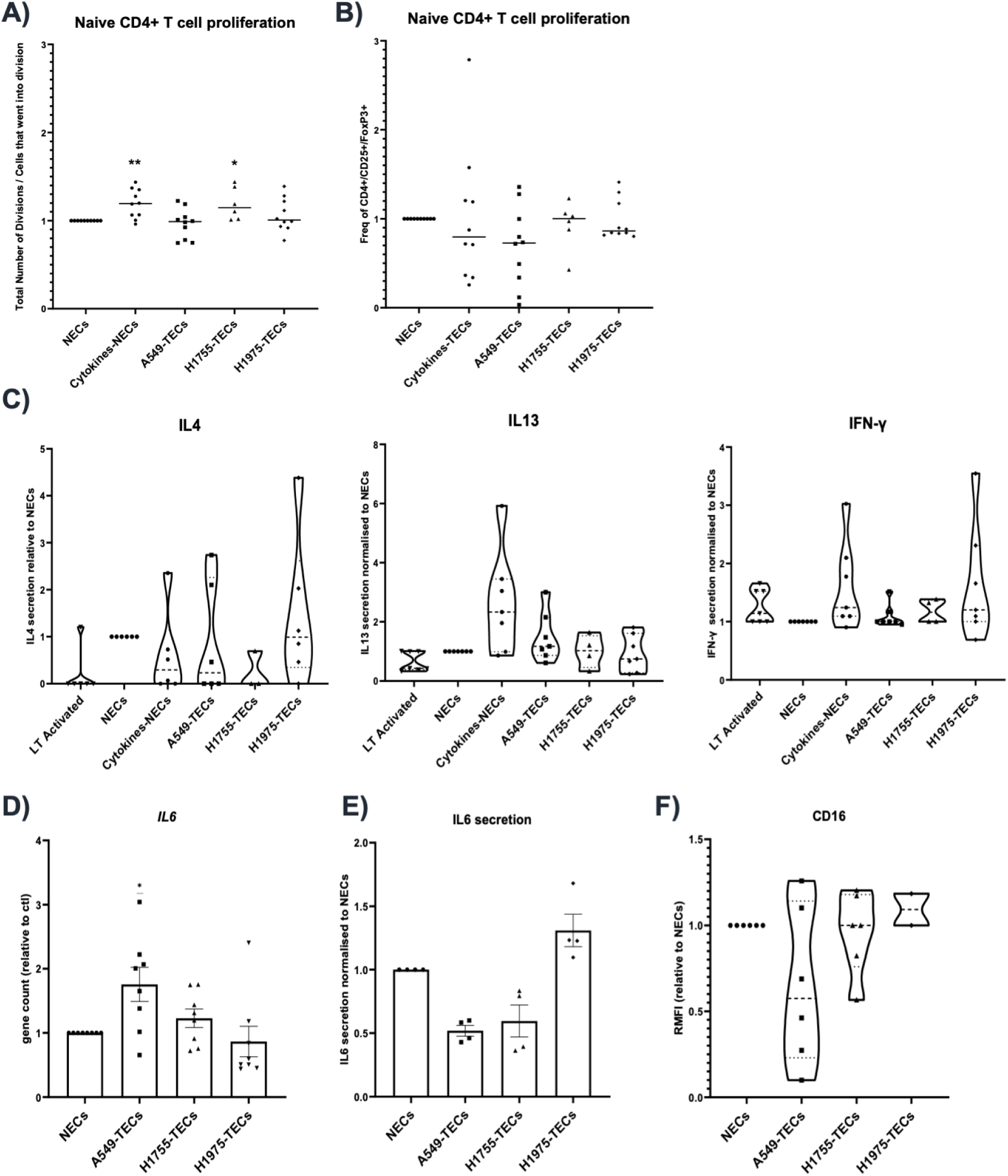
A) Proliferation of and B) Treg polarization of naive CD4+ T cells in cocultures with NECs or NSCLC-TECs and assessed with CFSE by flow cytometry. C) Cytokine secretion detected by Legendplex in naïve CD4 T cells/HUVEC cocultures. D) Gene count expression from the 3’RNA-seq for IL6 and E) IL-6 secretion by LegendPlex in NECs and NSCLC-TECs. F) Flow cytometry analysis for the M2-like polarization markers CD16 after the coculture between monocytes and NECs or NSCLC-TECs. Data are mean ± SEM, n > 4, *p< 0.05, **p < 0.01, Wilcoxon test compared to NECs.

**Supplementary Figure 5:**
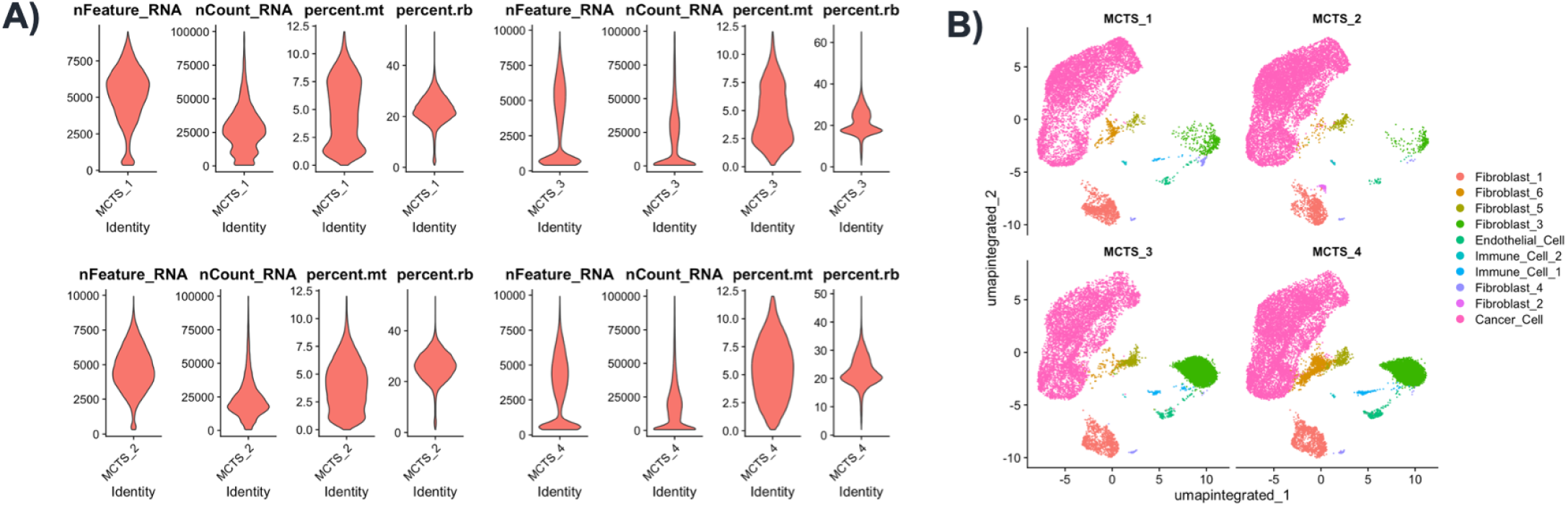
A) Violin plots of the quality control metrix with the number of Features, count and percentage of mitochondrial (mt) and ribosomal (Rb) genes. B) uMAP depicting cell clustering across all samples.

**Supplementary Figure 6:**
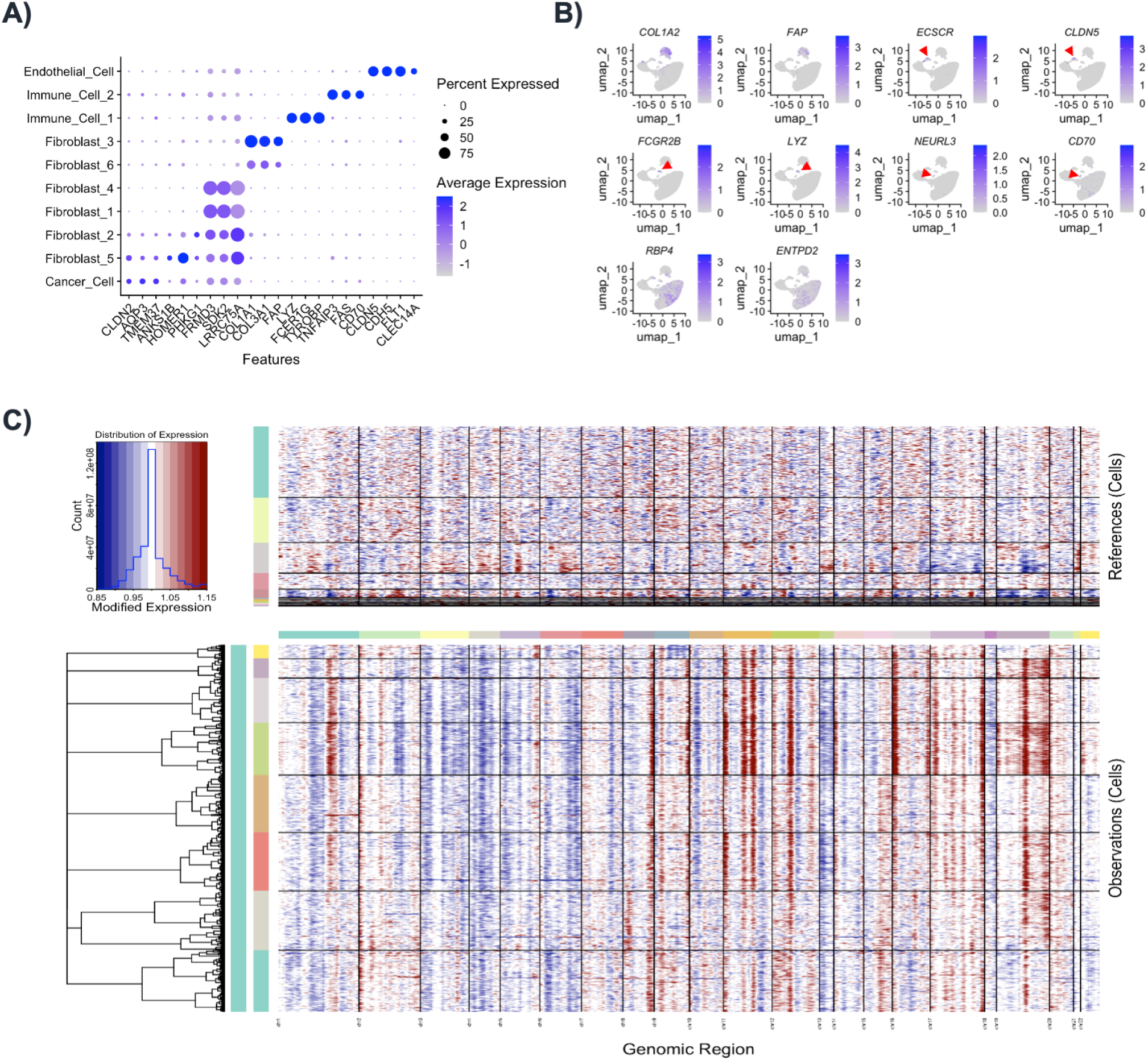
A) Dotplot showing key marker genes for the different MCTS clusters before merging the clusters belong to the same cell type. B) uMAP representation of canonical marker genes. C) Heatmap representing the chromosomal abnormalities in cells from the scRNA-seq of MCTS, as inferred by inferCNV. Reference cells at the top are endothelial, fibroblast and immune cells clusters, while the cancer cells are presented in the heatmap at the bottom and show several CNV (red, gain; blue, loss).

**Supplementary Figure 7:**
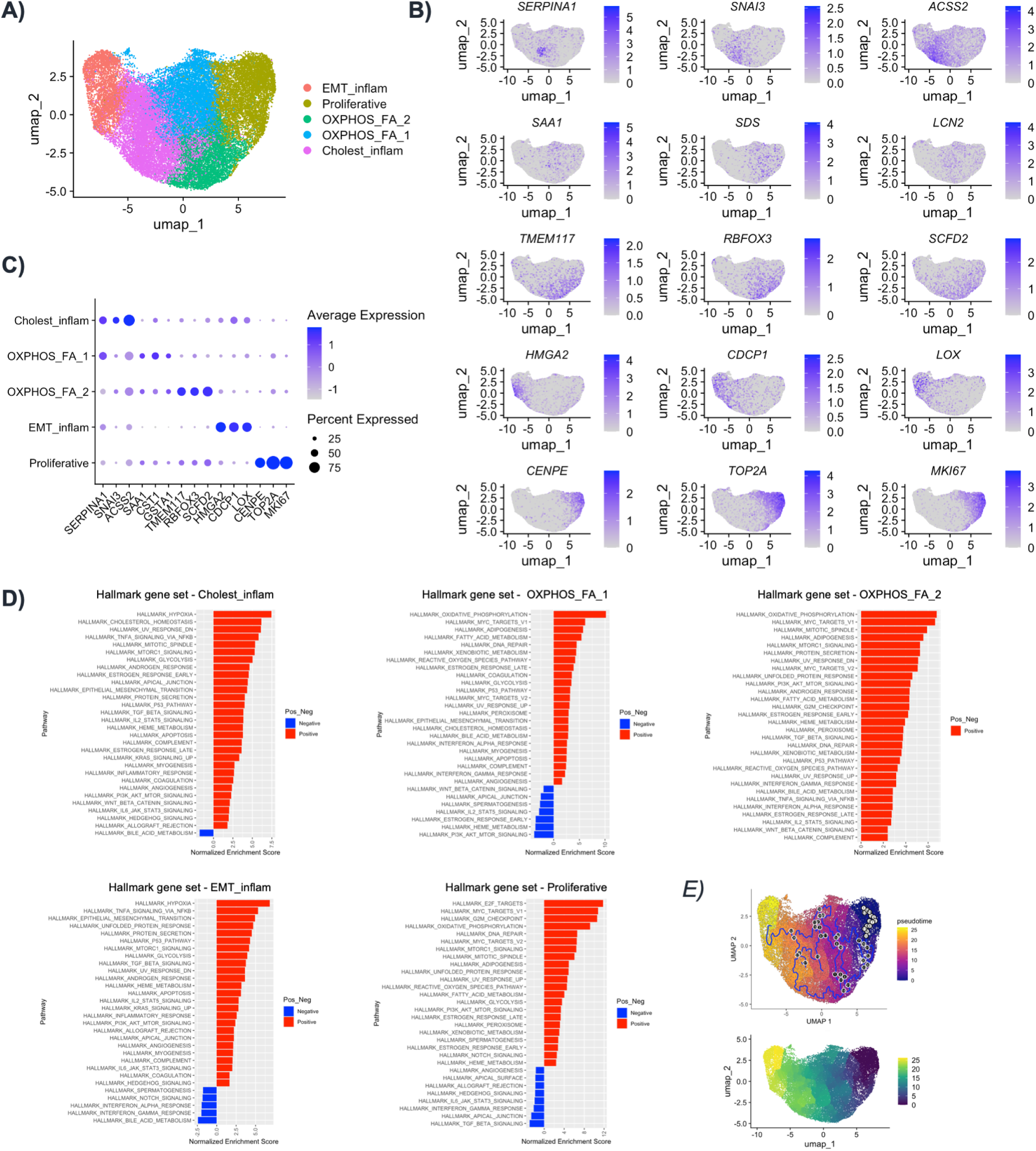
A) uMAP representation of the cancer cell subsets present within MCTS. B) uMAP representation and C) dot plot of the marker genes for each cancer cell subcluster. D) Gene set enrichment analysis for each cancer cell subcluster according to the Hallmark gene set. E) Trajectory analysis (top) and pseudotime uMAP representation of the cancer cell subset using Monocle.

**Supplementary Figure 8:**
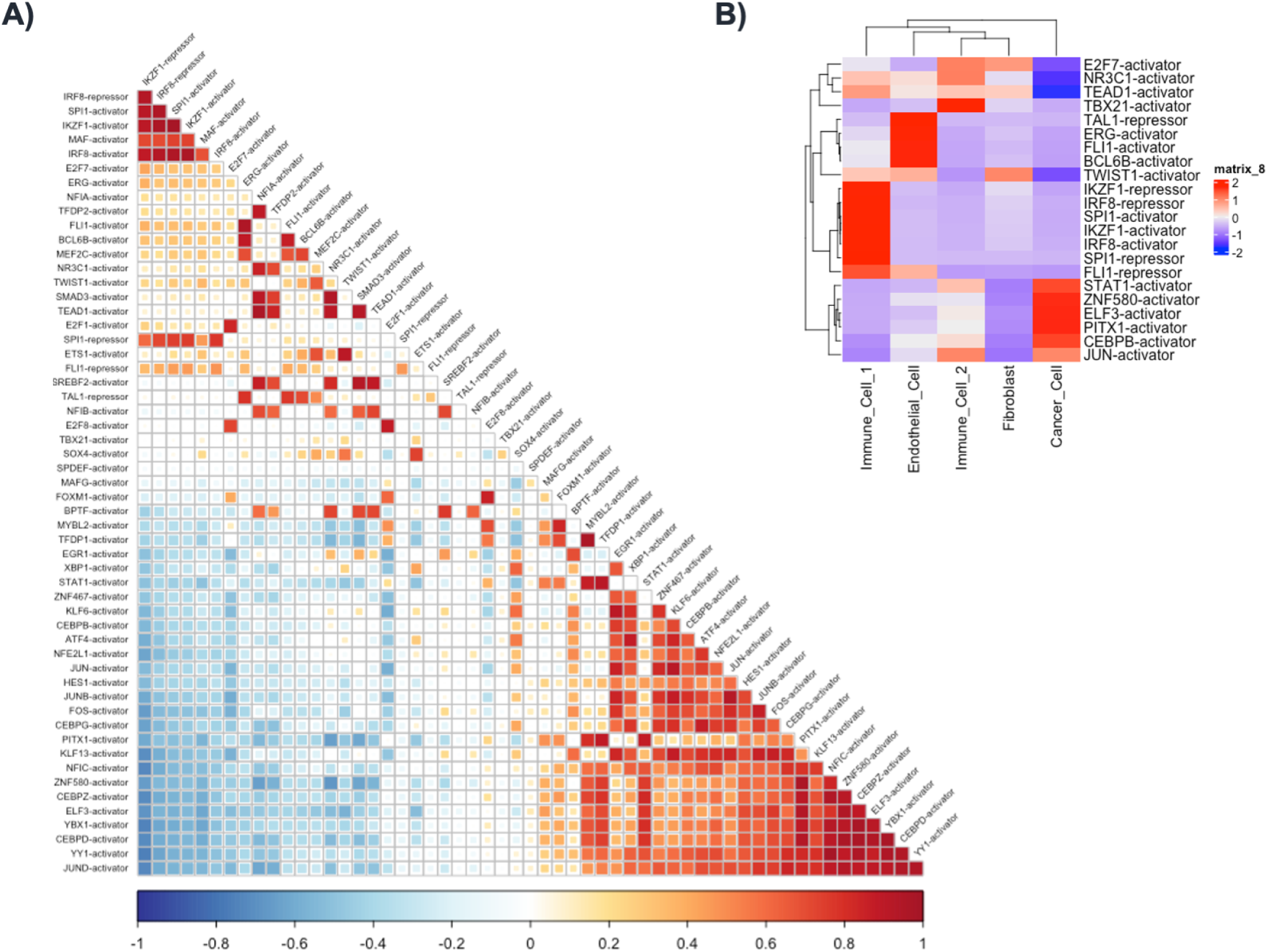
A) Correlation plot depicting co-regulated transcription factors. B) Heatmap of the most specific regulons per cell type.

**Supplementary Figure 9:**
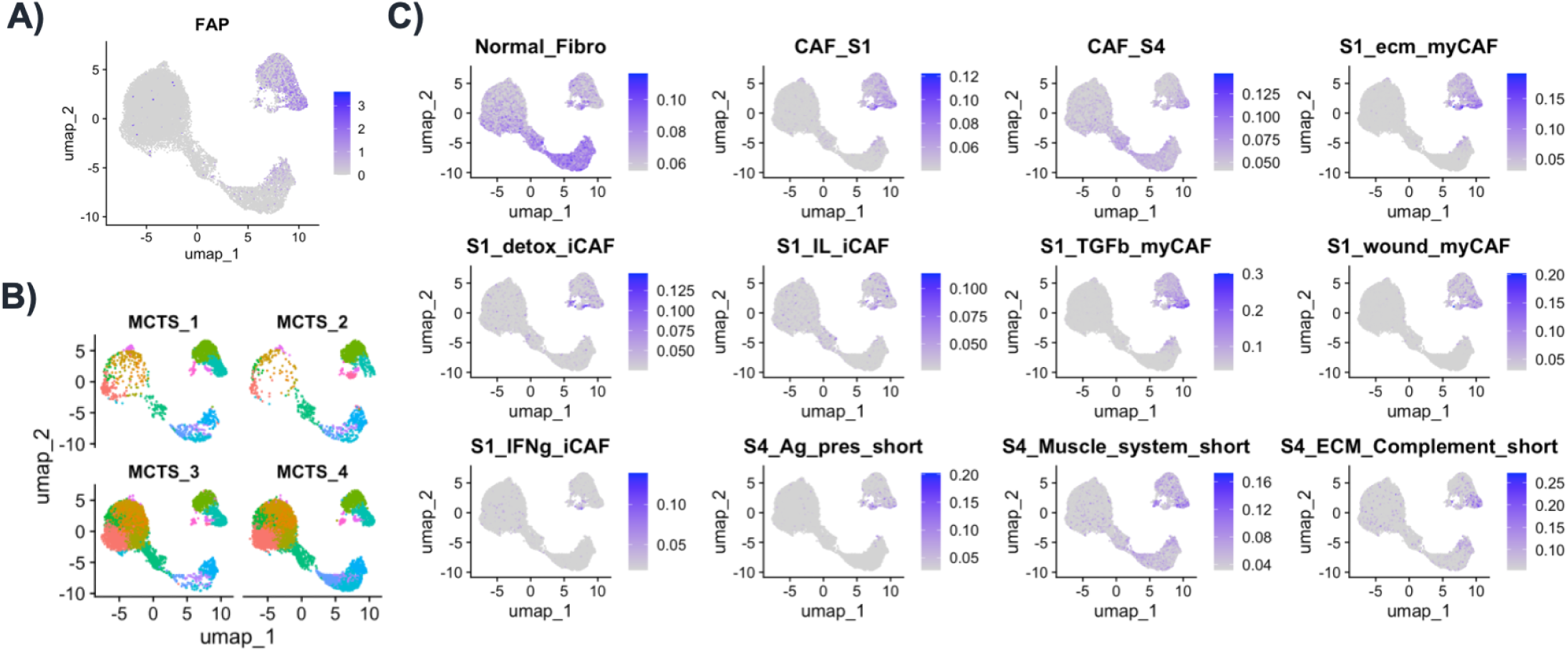
A) Feature plot showing the expression of the cancer-associated fibroblast (CAF) marker FAP across all cell type clusters from the MCTS. B) uMAP showing the fibroblast cluster with C) marker gene signatures for the major CAF subtypes identified previously ^40^.

**Supplementary Figure 10:**
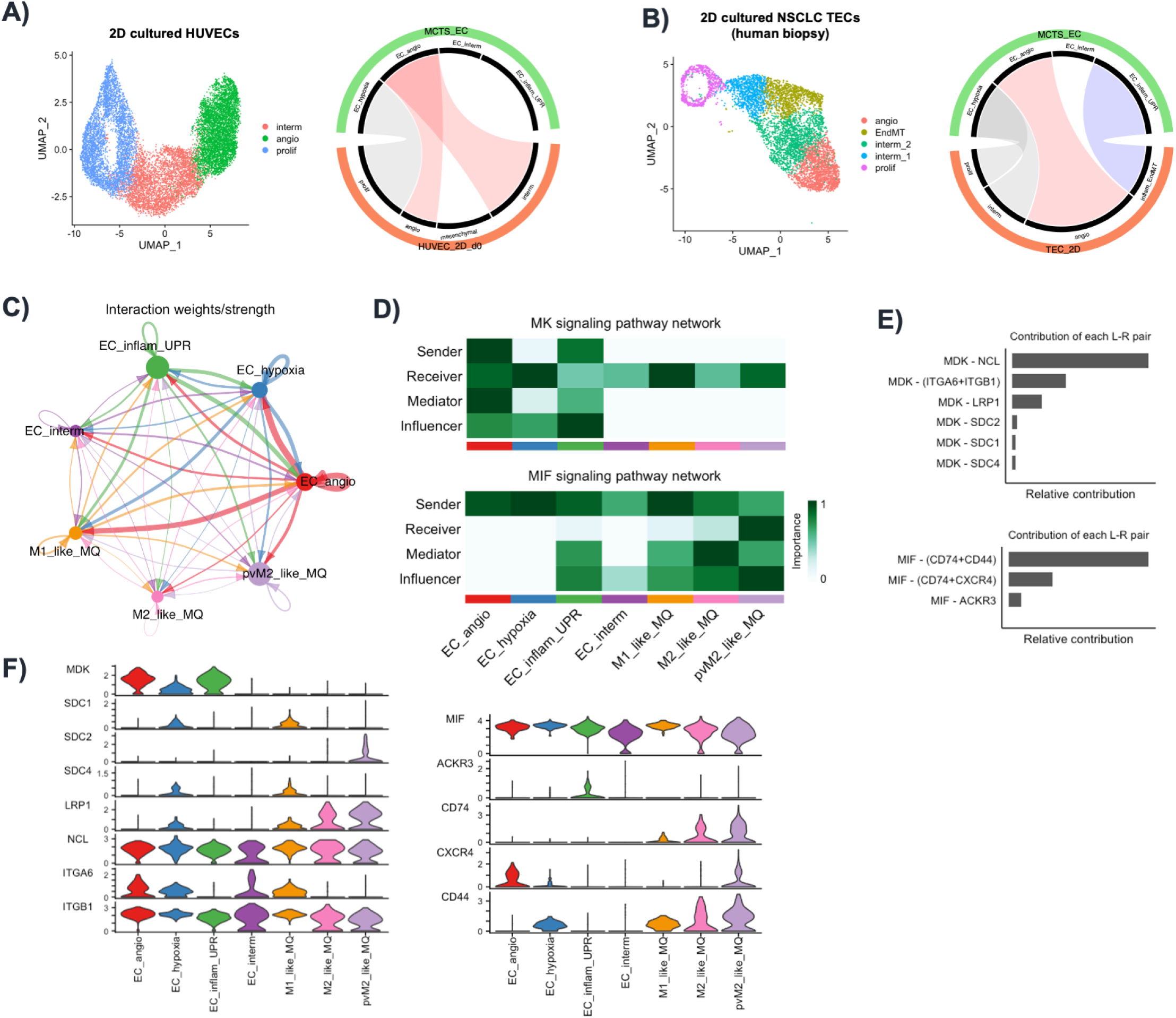
Circleplots showing the similarities between MCTS-EC subsets and A) 2D-cultured normal HUVECs, and B) TECs isolated from NSCLC human biopsies. The inflammatory MCTS-EC cluster was absent in 2D HUVECs, but present in 2D cultured TECs from patients however associated with mesenchymal features rather than inflammation. C) Chord diagrams showing the interaction weight between each considered clusters. Key regulators of the MK and MIF signaling pathway networks as depicted by D) heatmap for senders and receivers, E) contribution of each ligand-receptor pair, and F) violin plots.

